# Distinctive mechanisms of epilepsy-causing mutants discovered by measuring S4 movement in KCNQ2 channels

**DOI:** 10.1101/2022.01.19.476944

**Authors:** Michaela A. Edmond, Andy Hinojo-Perez, Xiaoan Wu, Marta E. Perez Rodriguez, Rene Barro-Soria

## Abstract

Neuronal KCNQ channels mediate the muscarine-regulated M-current, a key regulator of membrane excitability in the central and peripheral nervous systems. Mutations in KCNQ2 channels cause severe neurodevelopmental disorders, including epileptic encephalopathies. However, the impact that mutations have on channel function remain poorly defined, largely because of our limited understanding of the voltage sensing mechanisms that trigger channel gating. Here, we present measurements of voltage sensor movements in wt-KCNQ2 and channels bearing epilepsy-causing mutations using mutagenesis, cysteine accessibility, and voltage clamp fluorometry (VCF). Cysteine modification reveals that a stretch of 8-9 amino acids in the S4 become exposed upon opening of KCNQ2 channels. VCF shows that the voltage dependence and kinetics of S4 movement and channel opening/closing closely correlate, suggesting an activation scheme in which channel opening does not require multiple voltage-sensor movements. VCF and kinetic modeling reveal different mechanisms by which epilepsy-causing mutations affect KCNQ2 channel voltage-dependent gating. This study provides insight into KCNQ2 channel function, which will aid in uncovering the mechanisms underlying channelopathies.

## Introduction

Voltage-gated K^+^ (Kv) channels play a crucial role in regulating excitability and dysregulation of their function has been associated with several disorders, including cardiac arrhythmias, epilepsy and autism. Kv channels, including all members of the Kv7 family (Kv7.1–Kv7.5, also known as KCNQ as they are encoded by KCNQ1-5 genes (1–3)), are highly heterogenous and widely expressed in excitable cells where they regulate resting membrane potential, shape the firing and duration of action potentials, and control rhythmic events(4).

One of the major potassium currents throughout the central and peripheral nervous systems is the muscarine-regulated M-current. The M-current is mainly conducted by heterotetrameric combinations of KCNQ2/3 and KCNQ3/5 channel (5–8), but homotetrameric assemblies of channel subunits have also been shown to function as the M-current in neurons (8, 9). KCNQ are non-inactivating channels with slowly activating and deactivating kinetics and a negative voltage for half-activation(5, 6). These biophysical properties make KCNQ channels important regulators of neuronal excitability. For example, the peculiar lack of inactivation at voltages near the threshold for action potential initiation confers KCNQ channel’s dominant role in regulating membrane excitability as one of the main outward sustained currents. Thus, inhibition of the KCNQ channel lowers the action potential threshold, increase afterdepolarization(10), and slows excitatory postsynaptic potentials(11), resulting in a reduced adaptation and prolonged repetitive neuronal firing(12). Among the KCNQ family of proteins, KCNQ2 channels have received particular attention because mutations in this channel cause a variety of neurodevelopmental phenotypes(2, 13, 14), including epileptic encephalopathy (15–20), and more recently autism(21). Since KCNQ2 channels are central to physiological and pathophysiological events, it is important to understand the voltage-dependent mechanisms underlying channel opening to define its role in physiological control of neuronal excitability and to understand how specific variants alter KCNQ2 channel function.

The recently elucidated cryo-EM structure of human KCNQ2 channels(22) shows that, like canonical Kv channels(23), KCNQ2 contains six transmembrane helices (S1–S6) with cytosolic oriented N- and C- termini that form functional tetramers. The S5-S6 of the four subunits form a centrally located potassium selective pore that is flanked by the four voltage-sensing domains (S1-S4)(23). KCNQ2 has a domain-swapped tetrameric architecture, where the C-terminal end of the S6 segments form the gate(23, 24). The fourth transmembrane segment is assumed to be the voltage sensor (S4) since it contains several highly conserved positively charged amino acid residues. Although the S4 has been shown to move in response to changes in membrane voltages and functions as the voltage sensor in other Kv channels (25–28), this has not yet been directly demonstrated for KCNQ2 channels. Gating current recordings have not been resolved for KCNQ2 channels, likely due to low channel density within the membrane and/or the slow kinetic of activation compared to other Kv channels(29). Insight into S4 movement of KCNQ2 have been inferred by previous mutagenesis studies showing that charge neutralization of the arginine residues in S4 altered the voltage sensitivity of channel opening(30, 31). In addition, a disulfide crosslinking study showed that cysteine-substituted residues in the extracellular end of S4 crosslinked with a cysteine in S1 only in the closed state, further implying movement of S4(32). Although these studies provided insight into S4 rearrangements, they were unable to explain how S4 moves, nor were they able to offer a dynamic view of S4 motion during voltage-controlled gating of KCNQ2 channels.

Our understanding of the voltage-controlled activation mechanisms of KCNQ2 channels is limited, compared to other Kv channels like the related KCNQ1, whose physiological role in cardiac tissue has been extensively investigated(33). Kv channel opening can occur either after all four S4 have been activated(34), or alternatively through independent activation of each S4(35), as also reported for KCNQ1(36). Interestingly, pore opening of KCNQ1 channels can occur from two defined S4 conformations involving intermediate and fully activated S4 states(37–39). This activation scheme in which opening may occur from multiple S4 states, has provided a valuable framework to understand voltage-dependent gating of KCNQ1 with different accessory subunits, thereby allowing interpretation of its versatile physiology. Because we lack mechanistic understanding of voltage-dependent gating in neuronal KCNQ2 channels, it is not clear what mechanisms could be altered in the presence of disease-causing mutations. We here describe mechanisms underlying voltage sensor movement in KCNQ2 channels and propose a simple model of channel opening as it relates to epilepsy-associated mutations.

We provide an extensive exploration of positions where cysteine could be inserted into the S3-voltage sensor (S4) loop and S4 helix and used with both voltage clamp fluorometry (VCF)(25) and cysteine accessibility(26) to study S4 activation and its influence on disease causing mutations. Cysteine accessibility reveals that a stretch of 8-9 S4 residues becomes exposed upon opening of KCNQ2 channels. VCF shows that the voltage dependence and kinetics of S4 movement and channel opening/closing closely correlate. Our gating scheme suggests that multiple voltage sensor movements are not required to open KCNQ2 channels. VCF data and kinetic modelling shows that two epilepsy-causing mutations-R198Q and -R214W alter channel opening through two distinct mechanisms: R198Q shifts S4 movement and R214W changes the voltage sensor-to-pore coupling. These results provide critical information about KCNQ2 channel gating that will aid in future studies on KCNQ2 channelopathies.

## Results

### State-dependent external S4 modifications consistent with S4 as voltage sensor

The combination of cysteine-scanning mutagenesis and methanethiosulfonate (MTS) derivative modification is a powerful tool to study conformational changes in ion channel gating. This methodology assumes that upon covalent modification of substituted cysteines, functional changes in channel gating occur (Figure 1A). We test whether the fourth transmembrane domain (S4) in KCNQ2 channel functions as the voltage sensor by measuring the state-dependent accessibility changes of introduced cysteines in the S4 (or in the S3-S4 linker) of KCNQ2 channels (Figure 1A-C). We assess the state-dependent modification rate of substituted cysteines by plotting the membrane-impermeable thiol reagents (MTS)-induced change in current against the cumulative exposure to MTS reagents at either hyperpolarized (closed) or depolarized (open) voltages (See materials and methods section and voltage protocols in Figure 1B). As previously used to demonstrate that the S4 crosses the membrane during gating of the Shaker channel(26), we assume that changes in the state-dependent modification rate of substituted-cysteines by externally applied MTS compounds would indicate that some residues in S4 move (outward) across the membrane during channel activation.

**Figure 1.**
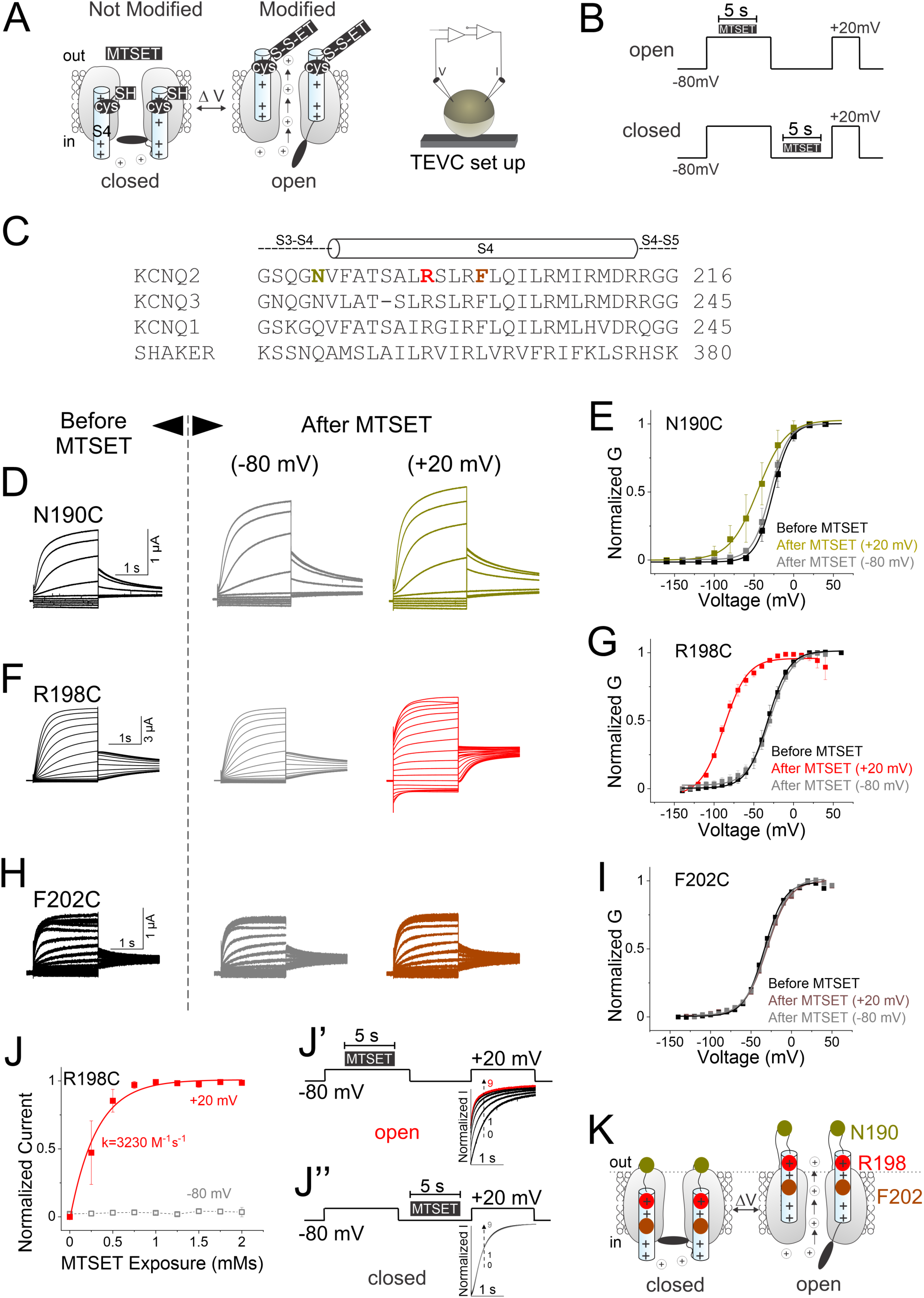
State-dependent modification of KCNQ2-R198C by external MTSET is consistent with outward S4 motion. (A) Cartoon of extracellular MTSET-modification of substituted cysteines (cys) of S4 during channel opening and a two-electrode voltage clamp (TEVC) setup. (B) Voltage/perfusion protocols in the open and closed states (see M & M). (C) Sequence alignment of the S4 region in Kv channels. (D, F, and H) Currents from oocytes expressing KCNQ2 -N190C (D), -R198C (F), and - F202C (H) channels in response to 20-mV voltage steps from –140 mV to +40 mV, (*Left*) before and after several 5-s applications of MTSET at (*Middle*) –80 mV and (*Right*) +20 mV. We repeat MTSET applications in between 25-s washouts for 8-12 cycles, as shown in (B). We used 1, 50, and 100 μM MTSET in (D), (F), and (H), respectively. (E, G and I) Normalized G(V) relations (lines from a Boltzmann fit) of recordings from panels(D), (F), and (H), respectively, before (black) and after MTSET application in the closed (-80-mV, gray) and open (+ 20-mV, colors) states (mean ± SEM, *n= 3-8*). (J) The rate of MTSET modification of R198C channels at +20-mV (red squares) or –80 mV (gray squares) was measured using the current amplitude at 400 ms after the start of the +20-mV voltage step (dashed line in J’ and J”, respectively). The current amplitude was plotted versus the cumulative MTSET exposure and fitted with an exponential. The fitted second-order rate constant in the open state protocol is shown in red. kopen = 3,230 ± 3.8 M^−^1 s^−^1 (*n = 3*). (J’) currents in response to a +20-mV voltage step during open-state 5-s MTSET application on R198C channels for the indicated voltage protocol. MTSET significantly increased the rate of activation. (J”) currents in response to a +20-mV voltage step during closed-state 5-s MTSET application on R198C channels for the indicated voltage protocol. MTSET application was repeated >8 times. (K) Cartoon representing the voltage-dependent cysteine accessibility data. MTSET modifies N190 in both the closed and open states. While F202 remains unmodified in both states (seemingly buried in the membrane), R198 becomes accessible only in the open state. Dashed line indicates the proposed outer lipid bilayer boundary.

In total, we made 8 cysteine mutants within the S4 (or in the S3-S4 linker) of KCNQ2 channels (Figure 1 and Figure 1-figure supplement 1). We express these mutants, one at a time, in *Xenopus* oocytes and use two-electrode voltage clamp to probe the external accessibility of the substituted cysteines to the MTS reagents (2-(ammonium)ethyl) methanethiosulfonate (MTSET) or sodium (2-sulphonatoethyl) methanethiosulfonate (MTSES) at both hyperpolarized (closed) and depolarized (open) voltages (Figure 1 and Figure 1-figure supplement 1). Overall, all cysteine mutants (N190C, A193C, S195C, A196C, R198C, S199C, L200C, F202C) have similar kinetics and steady-state conductance/voltage curve (G(V)) as wild-type-KCNQ2 channels (Figure 1D-I and Figure 1-figure supplement 1 and supplement Table 1).

MTSET and MTSES do not modify wt-KCNQ2 at either depolarized or hyperpolarized voltages (Figure 1-figure supplement 1A-C and supplement Table 1). However, external application of MTSET and MTSES modify the substituted-cysteine residue at position N190 in both the open and closed states (Figure 1D, E and Figure 1-figure supplement 1D, E). Modification of N190C channels quickly and irreversibly increases the current amplitude, speeds up the kinetic of activation, and changes the voltage dependence of activation in either closed or open N190C channels, albeit to distinct V_1/2_ values (as also shown earlier for HCN1 channels (40), Figure 1D, E), as if N190C is always accessible and exposed to the extracellular solution (Figure 1K, yellow). For R198C, external MTSET application in the open state (at +20-mV) irreversibly speeds up the kinetics of activation, increases the current amplitude, and left-shifts the G(V) relationship (Figure 1F-G, red). In contrast, when MTSET is applied in the closed state (at –80-mV), R198C channels are not modified (Figure 1F-G, gray). Since MTSET modifies residue R198C relatively fast at depolarized potentials (Figure 1J, J’, red) but not significantly at hyperpolarized potentials (Figure 1J, J’’, gray), that suggests that this residue is not accessible (i.e. is buried in the membrane) in the closed state (with S4 down) but becomes accessible in the open state (with S4 up, Figure 1K, red). Note that the perfusion system quickly delivers a 5-s pulse of MTS-reagents to the external surface of oocytes. This ensures perfusion of MTS-reagents at the indicated voltage as shown by the time course of solution exchange from 100 mM NaCl to 100 mM KCl (Figure 1-figure supplement 2). We find similar state-dependent modifications upon external MTSET perfusion for KCNQ2 channels with the S4-mutations A193C, S195C, A196C, S199C, and L200C (Figure 1-figure supplement 1F-O). External application of 0.1 mM MTSET (and even up to 1mM) showed no modification of channels with cysteines introduced further toward the C-terminus of the S4, such as F202C in either the open or closed states (Figure 1H, I). F202 is the first outermost N-terminal residue in the S4 segment to remain unmodified by external MTSET (Figure 1H-I). This result suggests that either F202C remains buried in the membrane during S4 activation (unmodified) even under conditions of high MTSET concentrations, long lasting and strong depolarization to + 20-mV (Figure 1K, brown), or alternatively that the modification does not significantly alter channel gating. Figure 1K, Figure 1-figure supplement 1P, and Figure 1-figure supplement 3 show cartoons summarizing a map of the voltage-dependent distribution of S4 residues in the resting and activated conformations inferred from the extracellular cysteine accessibility data.

### Tracking S4 movement of KCNQ2 channels using voltage-clamp fluorometry (VCF)

VCF, which relies on the physicochemical properties of fluorescent probes in different environments and on their short half-life once excited(41), allows simultaneous measurements of S4 movement (by fluorescence) and gate opening (by ionic current)(25) (Figure 2A). To track S4 movement in KCNQ2 channels using VCF, we first performed, one at a time, cysteine substitution of residues in the S3-S4 linker (Figure 2-figure supplement 1A). Compared to wild type KCNQ2 channels, cysteine mutations in this region only slightly changed the half-point of voltage-dependent activation (V_1/2_) (Figure 2- figure supplement 1A, B and supplement Table 1). Similar gating sensitivities have been shown for homologous cysteine mutations in KCNQ3 channels(42).

**Figure 2.**
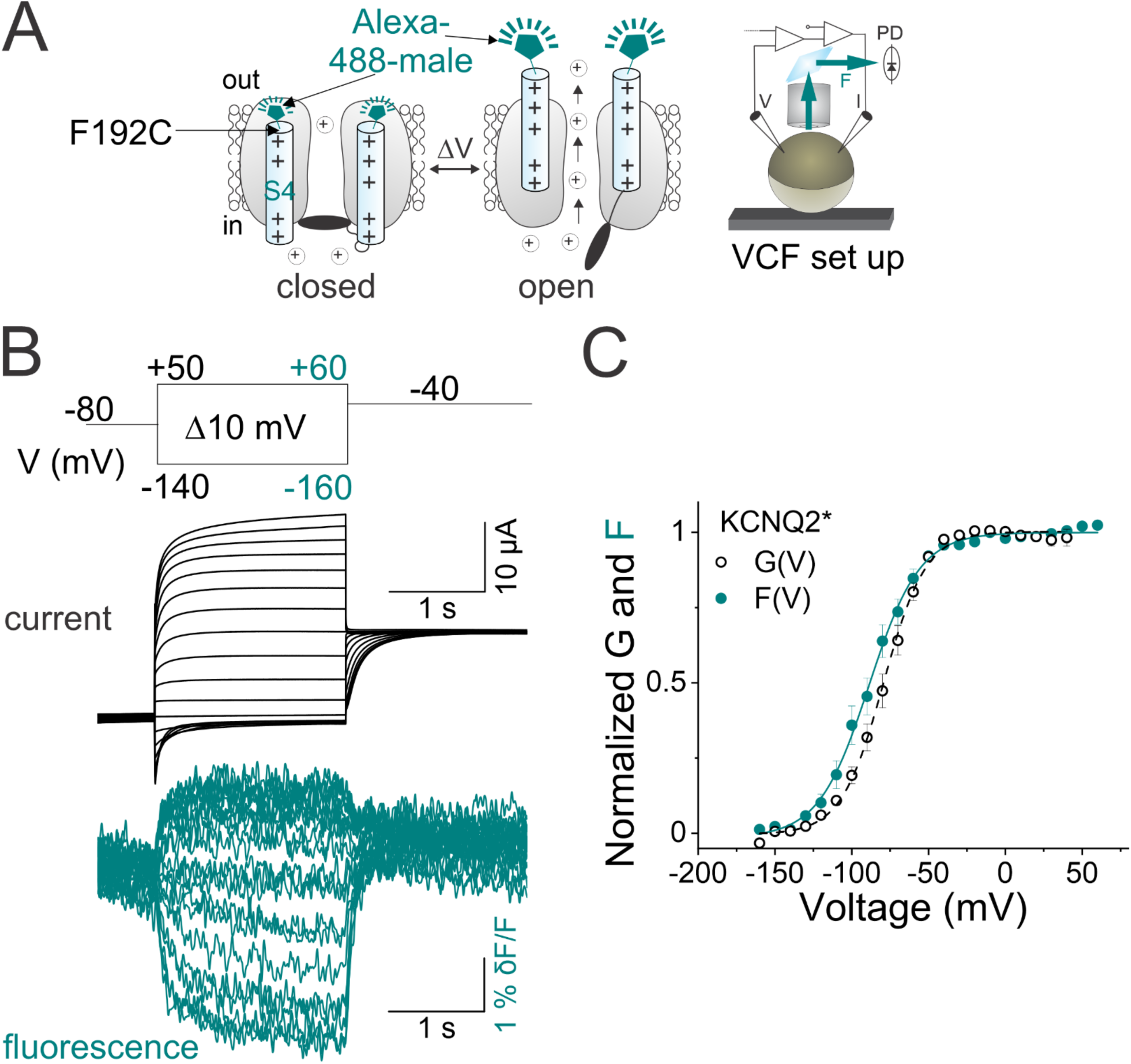
Labeled KCNQ2-F192C channels track S4 movement. (A) Cartoon representing the voltage-clamp fluorometry (VCF) technique. A cysteine is introduced at position 192 (close to the voltage sensor (S4)) and labeled with a fluorophore tethered to a maleimide group (Alexa-488-5 maleimide). Upon voltage changes, labeled- S4s move and the environment around the tethered fluorophore changes, altering fluorescence intensity. Both current (I) and fluorescence (F) are recorded simultaneously using a VCF set up. V, voltage; PD, photodiode photodetector. (B) Representative current (black) and fluorescence (cyan) traces from Alexa-488-labeled KCNQ2-F192C channels (KCNQ2*) for the indicated voltage protocol (*Top*). (C) Normalized G(V) (black circles and black dashed line from a Boltzmann fit) and F(V) (cyan circles and cyan dashed line from a Boltzmann fit) curves from KCNQ2* channels. The midpoints of activation for the fits are: G_1/2_ = –77.1 ± 2.7 mV, (*n = 8*) and F_1/2_ = –87.1 ± 3.9 mV, (*n=7*), supplement Table 1. Data are mean ± SEM; see M & M.

Of the five labeled KCNQ2 substituted cysteines, the mutant KCNQ2-F192C (henceforth called KCNQ2*) shows the most reliable and robust voltage-dependent fluorescence signals (maximum fluorescence change, dF/F ∼1%) that saturates well at negative and positive voltages, either labeled with Alexa488 5-maleimide (Figure 2B) or DyLight488-maleimide (Figure 2-figure supplement 1C, E). Compared to unlabeled KCNQ2* channels, the time courses of ionic currents of labeled KCNQ2* (labeled with either fluorophore) are similar, while their G(V)s are left-shifted by about ∼27mV (Figure 2-figure supplement 1D, E and supplement Table 1). Moreover, the time courses of fluorescence signal (labeled with either fluorophore) follow each other (Figure 2-figure supplement 1D, right panel). Importantly, the changes in fluorescence signal are likely caused by Alexa488-maleimide (or by Dylight488-maleimide) attached to F192C as oocytes expressing wild-type KCNQ2 channels treated with either fluorophore do not show a voltage-dependent fluorescence signal (Figure 2-figure supplement 1F, G). While both labeled KCNQ2* channels render similar fluorescence changes (Figure 2-figure supplement 1), we decided to use Alexa 488-5 maleimide to label KCNQ2* throughout the study.

Figure 2B shows the simultaneous measurement of ionic current (black) and fluorescence (cyan) from labeled KCNQ2* channels in response to a family of voltage steps (from –160 to + 60 mV) using VCF. The steady-state fluorescence/voltage curve, F(V), (assumed to reflect voltage sensor movement) closely tracks the steady-state conductance/voltage curve, G(V), (reflecting channel opening) in KCNQ2* channels (F(V) = –87.1 ± 3.9 mV, n= 7 and G(V) = –77.1 ± 2.7 mV, n= 9, Figure 2C and supplement Table 1). The fluorescence signals in Figure 2B have a non-linear voltage dependence and are much slower than the voltage changes per se, which suggests that the fluorescence changes are not electrochromic responses of the dye to voltage changes. Instead, these results suggest that the fluorescence changes around labeled F192C at the outer end of S4 are due to conformational changes of the S4 segment related to the opening and closing of the KCNQ2 channel gate.

Next, VCF was used to compare the time courses of fluorescence signals and ionic currents of KCNQ2* channels during both depolarization-induced activation and repolarization-induced deactivation (Figure 3). We use a pre-pulse to −120-mV to completely close the channel before stepping to test voltages (Figure 3A, E). The fluorescence signal decreases in response to the pre-pulse to −120-mV (Figure 3A, E, cyan arrow), indicating that not all voltage sensors are in their resting position at the −80-mV holding potential. We find that both ionic currents and fluorescence signals follow a double exponential time course in response to a family of voltage steps between −60 and +40-mV following the pre-pulse at −120-mV (Figure 3A, B). The fluorescence signal only slightly precedes the ionic current. While the voltage-dependence of the fast fluorescence component is smaller than the slow component, it shows clear differences between extreme voltages (Fig. 3C). Additionally, the time constant for the fast fluorescence component is smaller than time constant of the fast component of the current, whereas the time constants of the slow component of the fluorescence signal and ionic current closely follow each other at all voltages tested (Figure 3B, C). To estimate the contribution of the fast and slow components, we plot the fast and slow amplitudes of the fluorescence signal against voltage. At voltages more positive than 0 mV the amplitude of the fast fluorescence signal increases compared to the slow component so that the fluorescence signal is mainly due to the fast component (Figure 3D). In response to voltage steps between −80 and −180- mV to close the channels, both ionic currents and fluorescence signals follow a double exponential time course (Figure 3E, F). The ionic current overlaps fairly well with the fluorescence signal (Figure 3E-G). At increasingly negative voltages, the amplitude of the fast fluorescence signal increase compared to the slow component so that the fluorescence signal is predominantly due to the fast component (Figure 3H).

**Figure 3.**
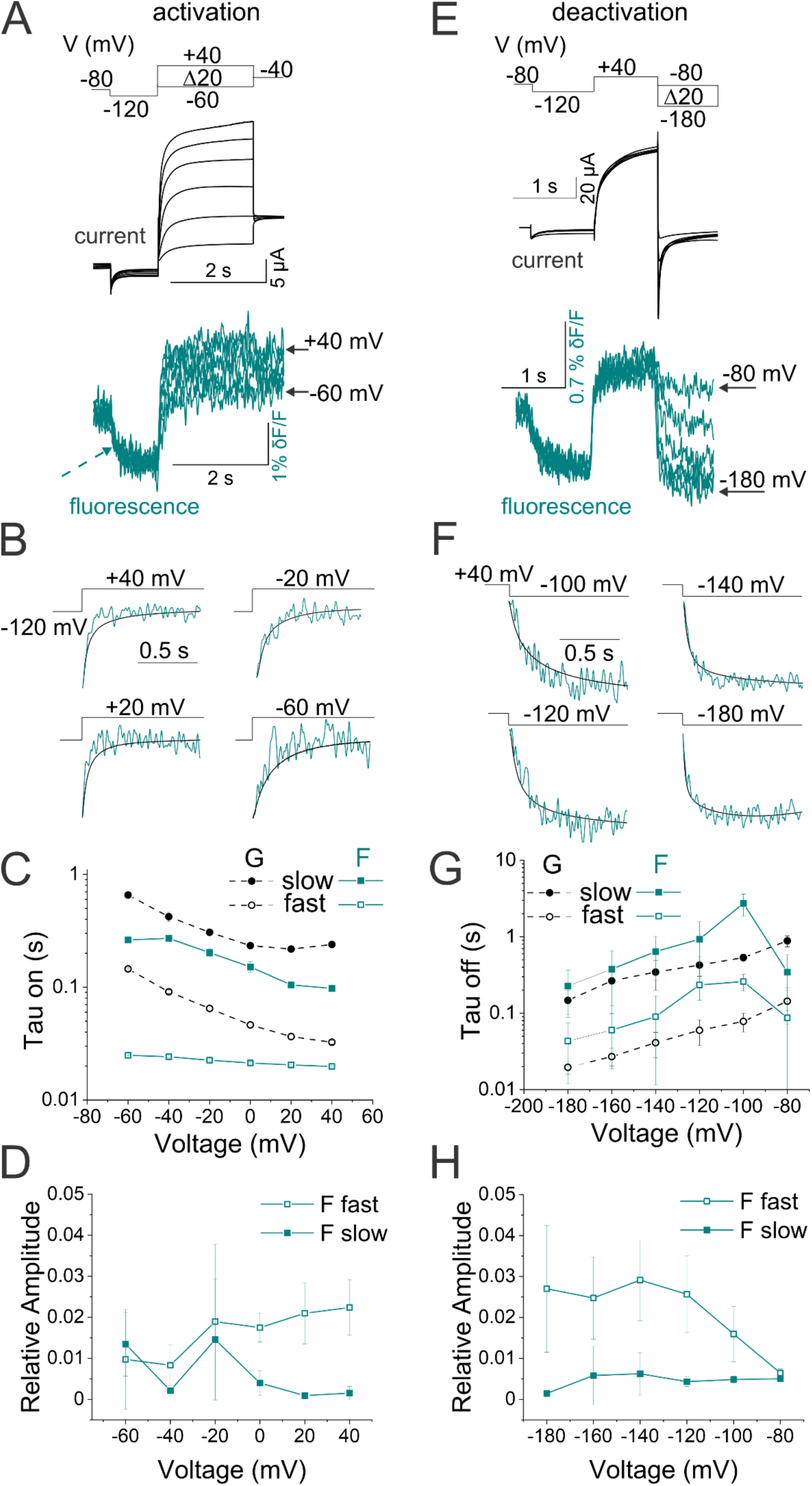
Fluorescence from KCNQ2* correlates with channel opening. (A-F) Representative current (black) and fluorescence (cyan) traces from KCNQ2* channels for the activation (A) and deactivation (E) voltage protocols (*Top*). In response to the prepulse to −120 mV, the fluorescence signal decreases (cyan dashed arrow), indicating that not all voltage sensors were in their resting position at the holding potential (−80 mV). Representative experiments showing (B) activation and (F) deactivation kinetics of current (black) and fluorescence (cyan) signals from KCNQ2* channels at different voltages as in (A) and (E), respectively. Note that the current and the fluorescence signals correlate, and they were fitted with double exponential curves. (C and G) Kinetic analysis of the voltage dependence of the conductance (black symbols) and fluorescence (cyan symbols) from (A-B) and (E-F), respectively. (D and H) Amplitudes of the fast (open squares) and slow (closed squares) fluorescent components from (A-C) and (E-G), respectively. Data in C, D, G, H (*n=4-7*) are mean ± SEM; see M & M

Collectively, these close correlations in time (Figure 3) and voltage dependence (Figure 2C) of fluorescence and current suggest that the environmental changes around labeled F192C at the outer end of S4 rendered fluorescence signals that seem to report on S4 motion associated with the opening and closing of the channel gate. These data also suggests that an individual voltage sensor movement might be sufficient to open the channel.

### S4 accessibility supports voltage-dependent motion of the S4 segment

We test whether the voltage-dependent fluorescence signals measured with VCF faithfully report on the S4 movement. We took advantage of the state dependent modification of A193C by MTSET (Figure 1-figure supplement 1F, G) to measure the rate of access to MTSET at different voltages as an independent assay (Figure 4). External MTSET modification speeds up the activation of A193C channels and increases the current amplitude (Figure 4B). While MTSET modifies A193C channels at voltages more positive than −100 mV (Figure 4B, C), MTSET, however, modifies A193C 5-fold faster at +20-mV than at −100-mV (Figure 4C and supplement Table 1). The modification rate for A193C approaches zero between −140 mV and −160-mV, as if A193C is inaccessible at those voltages (Figure 4D). The voltage dependence of modification rate by MTSET (modification rate/voltage curve or ‘mod. rate(V)’) follows the conductance/voltage curve G(V) for A193C channels (mod. rate(V) = – 72.7 ± 19.6 mV, n = 3-8 and G(V) = –66.4 ± 1.95 mV, n = 8, Figure 4D). The mod. rate(V) of A193C has similar voltage dependence as the F(V) of KCNQ2* channels (F(V) = –87.1 ± 3.9 mV, n = 7, Figure 4E, as if the fluorescence accurately reflects on S4 movement.

**Figure 4.**
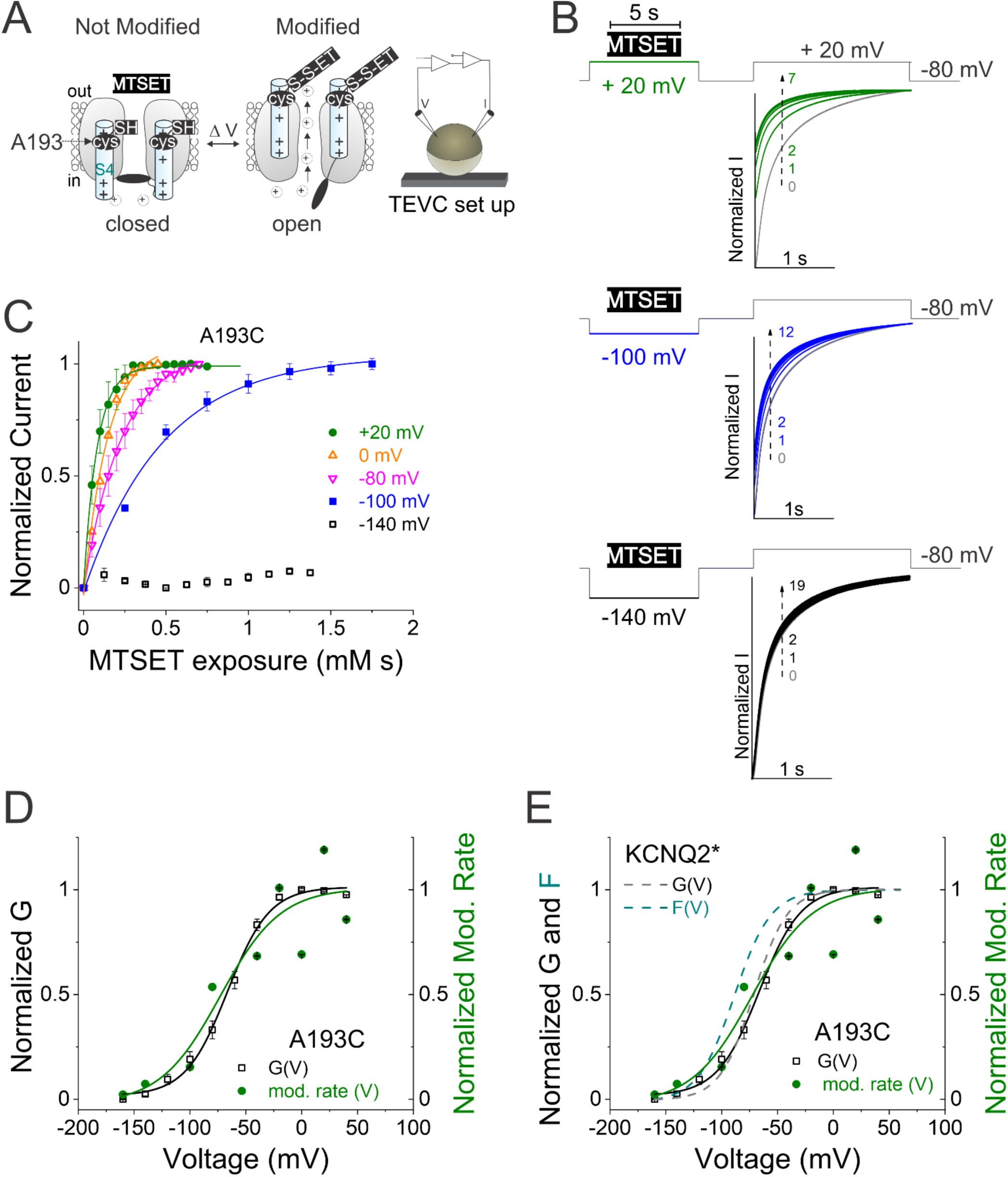
Accessibility of residue A193C supports voltage-dependent motion of S4 segment. (A) Cartoon representing extracellular cysteine accessibility of residue A193C as in Figure 1A. (B) Currents in response to +20-mV voltage steps before (gray trace 0) and during several 5-s applications of MTSET at +20-mV (green traces 1-7), –100-mV (blue traces 1-12), and –140-mV (black traces 1-19) on A193C channels for the indicated voltage protocol. We repeat MTSET applications (10 μM at +20- and –100-mV, and 20 μM at –140-mV) in between 25-s washouts as shown in each voltage protocol. (C) Normalized current of A193C during MTSET exposure at +20mV (green), 0mV (orange), -80mV (pink), -100mV (blue) and -140mV (black). (D and E) Normalized G(V) curve (squares and black line from a Boltzmann fit) of A193C channels and voltage dependence of the modification rate (mod. rate (V), green circles and green line from a Boltzmann fit) for MTSET to residue A193C. In (E), dashed lines represent ‘wt’ KCNQ2* G(V) (black) and F(V) (cyan) curves for comparison. The midpoints of activation for the fits are: A193C (mod. rate)1/2 = –72.7 ± 19.6 mV, (*n = 3-8)*; A193C G_1/2_ = –63.9 ± 1.4 mV, (*n = 11*). Data are mean ± SEM; see figure supplement 1 and Figure 2C for (wt) KCNQ2* G_1/2_ and F_1/2_, values.

### Disease-causing mutations differentially affect S4 and gate domains

Next, we took advantage of the epilepsy-causing mutations R198Q and R214W in KCNQ2 channels (43, 44) to further test if the voltage-dependent fluorescence changes around labeled F192C correspond to S4 motion. R198Q, which neutralizes the first gating charge of S4 in KCNQ2 channels (Figure 5A), was previously shown to shift the G(V) to more hyperpolarized potentials, to accelerate the kinetic of activation and to slow down kinetic of deactivation(43). We reason that if the fluorescence signal of the KCNQ2* channel observed in Figure 2 indeed reports on S4 movement, KCNQ2* bearing R198Q would similarly shift the F(V) curve to negative voltages and affect activation and deactivation kinetics. We find that the KCNQ2*-R198Q mutation causes a hyperpolarizing shift in the G(V) curve, slightly speeds up the kinetic of activation, and slows the kinetics of deactivation (Figure 5A-C and Figure 5-figure supplement 1A, B), as previously reported(43). VCF shows that KCNQ2*-R198Q channels exhibit fluorescence signals and ionic currents that continue to closely follow each other (Figure 5B and B’), further suggesting that the fluorescence change reports on S4 movement coupled to channel opening. KCNQ2*-R198Q channels exhibit fluorescence signals that are shifted to negative voltages (ΔF_1/2_ = − 35.1 ± 1.4 mV) similar to its negatively shifted G(V) curve (ΔG_1/2_ = − 30.1 ± 1.3 mV) (Figure 5C). These results further suggest that the fluorescence signal from F192C channels tracks the voltage-sensing rearrangement of S4 that controls channel opening, assuming that the R198Q mutation directly affects movement of the S4.

**Figure 5.**
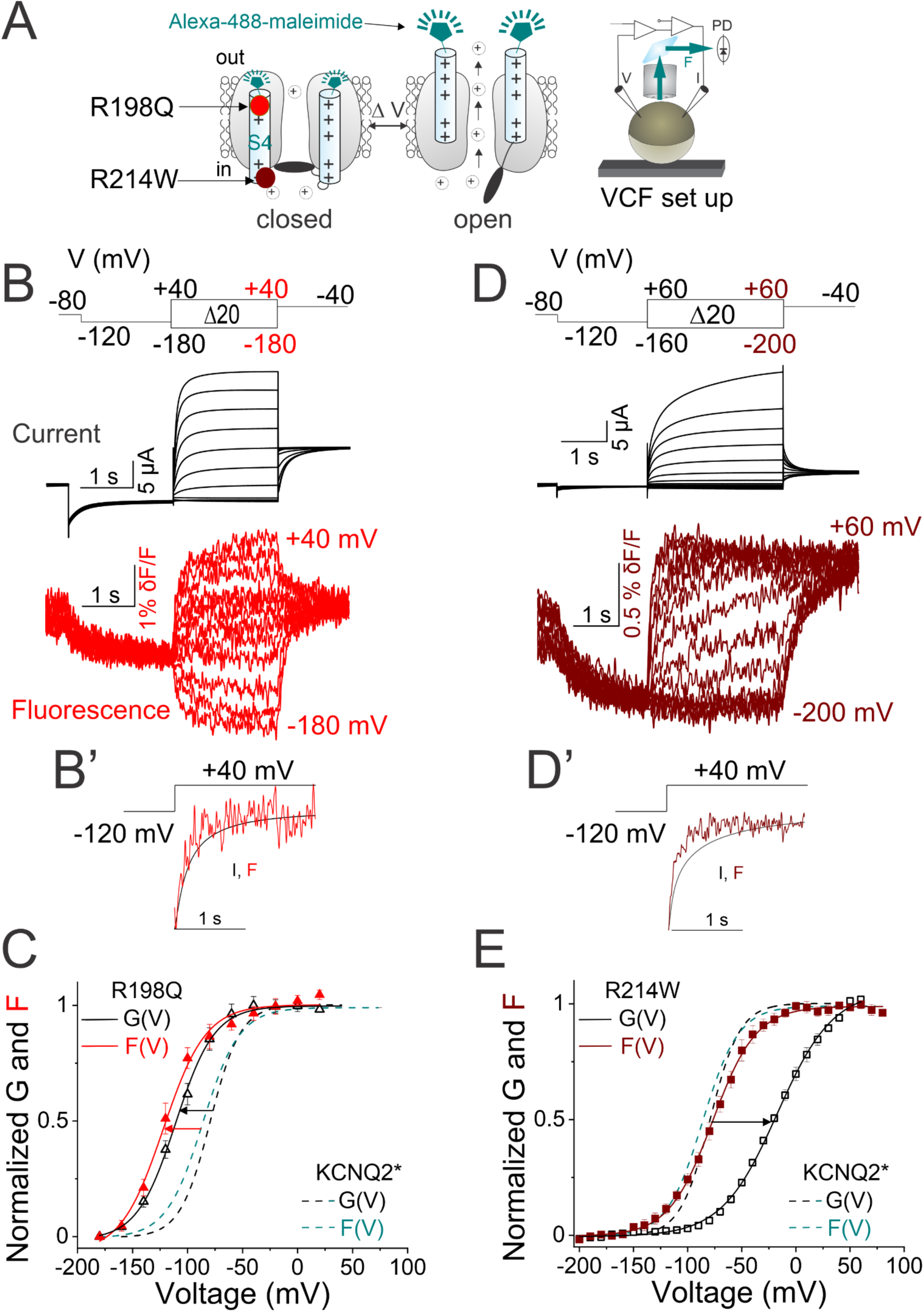
Disease-causing mutations in KCNQ2 channels differentially affect S4 and gate domains. (A) Cartoon representing the voltage-clamp fluorometry (VCF) technique as in Figure 2A. The localization of the two epilepsy-causing mutations- R198Q (red) and R214W (maroon) are shown. (B) Representative current (black) and fluorescence (red) traces from KCNQ2*-R198Q channels for the indicated voltage protocol (*Top*). (B’) Comparison of activation kinetics of current (black) and fluorescence (red) signals from KCNQ2*-R198Q channels in response to the voltage protocol shown. (C) Normalized G(V) (black triangles and black solid line from a Boltzmann fit) and F(V) (red triangles and red solid line from a Boltzmann fit) curves from KCNQ2*-R198Q. (D) Representative current (black) and fluorescence (maroon) traces from KCNQ2*-R214W channels for the indicated voltage protocol (top). (D’) Comparison of activation kinetics of current (black) and fluorescence (maroon) signals from KCNQ2*- R214W channels in response to the voltage protocol shown. (E) Normalized G(V) (black squares and black solid line from a Boltzmann fit) and F(V) (maroon squares and maroon solid line from a Boltzmann fit) curves from KCNQ2*- R214W. (C and E) Dashed lines represent ‘wt’ KCNQ2* G(V) (black) and F(V) (cyan) curves for comparison. The same color code for the two KCNQ2 mutations is shown throughout the figure. The midpoints of activation (G_1/2_ and F_1/2_) for the fits are shown in supplement Table 1. Data are mean ± SEM, (n = 4- 10).

In contrast to R198Q, the R214W mutation was previously reported to shift the G(V) relationship to more depolarized voltages(44). Compared to KCNQ2*, KCNQ*-R214W channels display a rightward shifted G(V) curve (ΔG_1/2_ = + 60-mV ± 1.8 mV, Figure 5E), and have a slower kinetics of current activation and faster kinetics of current deactivation (Figure 5-figure supplement 1C, D), as previously reported for R214W channels(44). VCF shows that in R214W channels, the time course of fluorescence signal precedes the time course of ionic current (Figure 5D and D’). Interestingly, the F(V) curve of R214W, which is + 10-mV shifted compared to the F(V) curve of KCNQ2* channels, is markedly left-shifted compared to its G(V) curve (Figure 5E). These results indicate that R214W seems to mainly affect the S6 gate and not directly the S4 segment. The separation between F(V) and G(V) suggests that, like the uncoupling mutation F351A in KCNQ1 channels(36, 37, 45) or the ILT mutation in the Shaker K^+^ channels(46), R214W dissociates voltage sensor movement from channel opening. Collectively, our results on the R198Q and R214W mutations provide additional support that the fluorescence signals observed from KCNQ2-F192C labeled with Alexa-488-maleimide are indeed reporting on voltage-gated conformational changes in the S4 preceding channel opening and not on voltage-gated conformational changes in other domains of the channel, such as gate opening. Furthermore, we hypothesize that since R214W is in the loop connecting S4 to the S4-5 linker (not within the voltage sensor, Figure 5-figure supplement 2A), it will most likely affect the activation gating without directly affecting S4 movement.

### Kinetic Model for KCNQ2 channel gating

We use the two protocols in Figure 3 to estimate the rates and voltage dependence of the S4 activation and deactivation transitions (Figure 6B-E). Upon depolarization, a first step (with rate *α*) involves S4 movement from a resting to an activated state. At more positive voltages, a second rearrangement of S4 occurs (with rate *γ*) to open the channel (Figure 6A). Upon hyperpolarization, the S4 rearranges back during channel closing (with rate *δ*), then it moves back to its resting state (with rate *β*) (Figure 6A). The rates *α* and *γ* are assessed by the time constant *τ_α_* and *τ_γ_* of the fast and slow (depolarization-induced activation) fluorescence components, respectively, in response to the indicated activation voltage steps (Figure 3A). *τ_α_* and *τ_γ_* approximate 1/*α* and 1/*γ* for sustained depolarizations rendering an effective gating charge z*α* = 0.43 ± 0.05 e0 and z*γ*= 0.48 ± 0.03 e0, respectively (Figure 6B, C). Likewise, the rates *δ* and *β* are assessed by the time constant *τ_δ_* and *τ_β_* of the fast and slow (hyperpolarization-induced deactivation) fluorescence components, respectively, in response to the indicated deactivation voltage steps (Figure 3D). *τ_δ_* and *τ_β_* approximate 1/*δ* and 1/*β* for sustained hyperpolarization rendering an effective gating charge z*δ* = 0.45 ± 0.06 e0 and z*β* = 1.18 ± 0.04 e0, respectively (Figure 6D, E).

**Figure 6.**
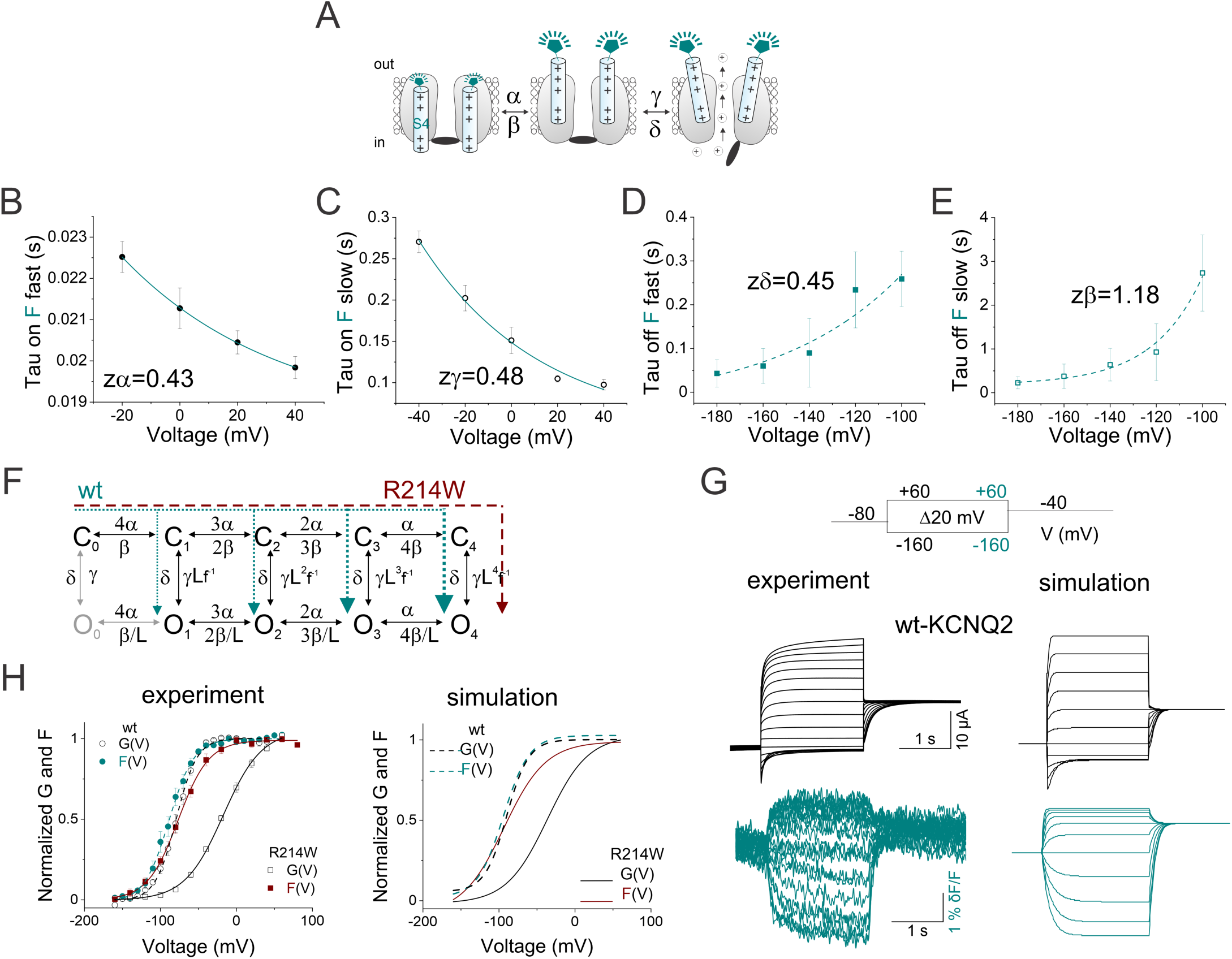
Model for KCNQ2 channel gating. (A) Cartoon representing a simplified model of KCNQ2-S4 activation and deactivation transitions with their respective rate constants. (B-E) Estimates of rate constants and voltage dependence of S4 (B and C) activation and (D and E) deactivation transitions of KCNQ2 channels from the VCF voltage protocols shown in Figure 3A and 3*B*, respectively. Voltage dependence of the time constant *τ* (which approximate (B) 1/*α*, (C) 1/*γ*, (D) 1/*δ* and (E) 1/*β*) were determined by fitting fluorescence traces from Figure 3 with a bi-exponential and using the fast and slow time constants for (*τ_α_*, *τ_δ_*) and (*τ_γ_*, *τ_β_*), respectively. Rate constants in (B-E) were of the form k_i_=k_o_ x exp(-z_i_(V)/*k*T), where k_i_ and z_i_ were determined from data in Figures 3 and 6. R, T and F have their usual thermodynamic meaning. Data in B-E are mean ± SEM, (*n=4-7*). (F) A ten-state gating scheme for KCNQ2 channels with the rates between transitions. The S4 movement is represented by horizontal transitions and closed/opening by vertical transitions. (G) Simulation of current (black) and fluorescence (cyan) for ‘wt’ channels using the protocol (*Top*). All parameters in the model are determined by the estimates of the rate constants and their voltage dependences from (B-E, see all parameters in supplement Table 2). Current and fluorescence were simulated using Berkeley Madonna (Berkeley, CA, USA). (H) Simulated G(V) and F(V) curves for ‘wt’ KCNQ2* (dashed lines) and R214W (solid lines) from simulation in (G). For comparison, experimental VCF data from Figure 2 are shown in (G) and (H). The cyan and maroon arrows in (F) denotes the likely transition pathway for the ‘wt’ and R214W channels, respectively.

To gain insight into the gating mechanisms of KCNQ2 channels, we use a Markov allosteric kinetic model (Figure 6F) to simulate current and fluorescence data from KCNQ2-F192C during channel gating, similar to the activation scheme initially proposed for KCNQ1 channels(36). In this model, voltage sensor (S4) movements between resting and activated conformations are represented by horizontal transitions and channel closing/opening by vertical transitions (Figure 6F). The opening does not require that all voltage sensors (S4) have moved. The more S4 moves, the stronger the probability of transitioning to the open state, as represented by the parameters (L, f) controlling the opening rate. We find no experimental evidence supporting a constitutive open state (O_0_ in Figure 6F, gray) as channels seem to fully close at voltages more negative than −140 mV (Figure 2).

We simulate KCNQ2 gating using the calculated rate constants and voltage dependences from Figure 6B-E. The calculated total gating charge moved for KCNQ2 channels is 7.37 *e*_0_/channel [= 4 x (z*α* + z*β*) + (z*γ* + z*δ*)]. Although the protocols in Figures 3 and 6B-E do not fully isolate the different rate constants, the parameters obtained are sufficient for our model. With this gating scheme (Figure 6F), we can qualitatively describe the 1:1 relationship between the time courses of fluorescence and ionic currents of KCNQ2 (Figure 6G) and simulate the close correlation of G(V) and F(V) curves from KCNQ2 (Figure 6H). By decreasing the opening transition (L in Figure 6F) relative to wt KCNQ2 channels, the model also describes the clear separation between F(V) and G(V) curves observed in mutated KCNQ2*-R214W channels (Figure 6H), under the assumption that R214W changes the voltage sensing domain-pore domain (VSD-PD) coupling such that it prevents opening before multiple S4 have activated (Figure 6F, dashed maroon arrow). Additionally, data from the R198Q mutation can also be simulated by shifting the voltage dependence to negative voltages. This model suggests that for wt-KCNQ2 channels, all four voltage-sensor movements are not required before gate opening. In contrast, R214W seems to require multiple transitions of voltage sensors between closed states before channel opening. This gating scheme represents a starting point to understand the voltage-sensing mechanisms that are coupled to KCNQ2 activation as well as the functional effects of disease-causing mutations.

## Discussion

In this paper, we provide functional data that support the hypothesis that the fourth transmembrane segment (S4) in KCNQ2 channels acts as the voltage sensor that promotes channel opening. Our fluorescence measurements show a close correspondence between the voltage sensor movement and channel opening in KCNQ2 channels as both voltage dependence and the time courses of fluorescence and ionic current closely correlate. VCF, mutagenesis, and kinetic modeling suggest that in KCNQ2 channels multiple voltage-sensor movements are not required to open the gate. We find that two epilepsy mutants cause shifts in voltage dependence of channel opening by two different mechanisms: R198Q shifts S4 movement and R214W changes the VSD-PD coupling such that channel opening only occurs after all voltage sensors have moved. Because KCNQ2 channels play a pivotal role in controlling neuronal excitability, our findings uncovering the dynamics and state-dependent molecular rearrangements that lead to channel gating, will be central to understanding how channelopathies alter neuronal function. Understanding how mutations affect channel activity through different mechanisms can lead to better ways to correct these mutational defects.

Using a state-dependent cysteine modification approach, we map the extracellular boundaries of S4 residues during membrane depolarization. Our cysteine accessibility data suggests that a stretch of 8-9 amino acids (193 to 200-201), about half of the 17-19 residues forming the S4(22), move from a membrane-buried position in the resting state to the extracellular solution during activation gating. A study using mutagenesis and disulfide crosslinking of substituted cysteines or metal-ion bridge experiments, inferred putative closed-resting states of S4 of KCNQ2 channels(32). In this study, the first and second positively charged residues of S4 (R198 and R201) were assumed to interact with the first and second counter-charge residues (E130 and E140) in the S2 segment. This arrangement positioned the gating charge transfer center in S2 (F137) in between R198 and R201 in what was assumed to be a deep closed-resting state of S4 (figure supplement 5A). More recently, the cryo-EM structure of KCNQ2 channels(22) (PDB: 7CR0) revealed the snapshot of the channel in its activated (S4 up) state and the pore in the closed state. This structure shows that R198 and R201 had now moved about three helical turns outward (upward) from F137 into a position close to or within the extracellular space(22) (Figure 1-figure supplement 3B). These rearrangements are in line with our cysteine accessibility data in which residues N-terminal to residue F202 become exposed in the activated state of S4 at strong depolarizations (Figure 1-figure supplement 3C, D).

This relatively large outward motion of S4 is consistent with our estimated total gating charge of 7.37 *e0 per* channel moved during KCNQ2 activation gating. Compared to the charges moved during gating of the Shaker K^+^ channel (12 to 14 elementary charges per channel(27, 28, 47, 48)), our estimates seem low. However, compared to the 7 arginines present in S4 of Shaker, the S4 of KCNQ2 channels has only 5-6 arginines, with the third one (R3-like) being a glutamine conserved across KCNQ channels. Interestingly, in Shaker channels, R3 seems to contribute around 1 *e0* per subunit (4 *e0*/channel)(28), which may explain why in KCNQ2 channels (with R3 substituted by glutamine) the calculated equivalent electronic charges moved across the membrane is less than the value of Shaker channels. Compared to the total charges moved during gating of related KCNQ1 channel (4.13 *e0* per channel)(49), the 7.37 e0 per KCNQ2 channel activation seems slightly bigger. However, in contrast to KCNQ2 channels, KCNQ1 only has 3 arginines in S4, with the third one (R5- like) being a histidine. Overall, our data suggests that S4 of KCNQ2 channels undergoes similar movement to that of KCNQ1 and Shaker K^+^ channels.

We found that the fluorescence signals from labeled F192C track conformational changes of S4 in KCNQ2 channels. VCF shows that both the steady-state voltage dependence of S4 transitions and the kinetics closely follow those of ionic currents, which concurrently had virtually no delay and exhibited no sigmoidal time course. These findings indicate that S4 does not necessarily require independent conformational changes in all four KCNQ2 subunits before channel opening as shown for classical voltage-gated K^+^ channel from the squid giant axon(50). Instead, these close correlations in time and voltage dependence of fluorescence and current support a gating scheme in which either the S4s move concertedly to open the channel, or individual voltage sensor movement might be sufficient for pore opening similar to that proposed for KCNQ1 (without KCNE1) channels(36, 51). Based on our KCNQ2 data, we cannot distinguish between concerted or individual S4 movement, which will require additional experiments that are out of the scope of this work. Using VCF on linked concatemers of KCNQ1 subunits, it was shown that all four voltage sensors move independently, and channel opening can proceed before all voltage sensors have moved(51). Subsequent studies on KCNQ1 refined this initial model and further showed that the S4 can adopt resting- and intermediate-states at negative voltages before reaching a fully activated state at depolarized voltages, and that pore opening can occur from either S4 states(37, 38). Additionally, these functional distinctive S4 configurations changed the ion permeation properties of KCNQ1 pore since the intermediate state showed a higher Rb^+^/K^+^ permeability ratio than the fully activated S4 state(37). The same study proposed that because KCNE1 reduced the Rb^+^/K^+^ permeability ratio of KCNQ1, KCNE1 bypassed the intermediate S4 state and forced KCNQ1 channels to open only from the fully activated S4 state. Here, we explored whether KCNQ2 channels also change their ion-permeation properties to get insight into pore conformations. We reason that if KCNQ2 channel opening also occurs with S4 in the intermediate state, the Rb^+^/K^+^ permeability ratio would be lower in the mutant R214W that dissociates voltage sensor movement from channel opening (Figure 6F, dashed maroon arrow) than in the wt- KCNQ2 channel, much like the effect of KCNE1 on KCNQ1 pore. However, we find very similar Rb^+^/K^+^ ratios for both wt (0.52 ± 0.016) and the R214W mutant (0.51 ± 0.024) (Figure 6-figure supplement 1A, B), as if R214W did not alter the pore conformation relative to the wt KCNQ2 channel. In contrast to KCNQ1(37), we see no experimental evidence of an intermediate-open state in KCNQ2 channel. Our results indicate that while the overall voltage-dependent gating mechanisms of KCNQ2 is qualitatively similar to that of KCNQ1(51), subtle differences exist (such as the experimental lack of an intermediate-open state in KCNQ2), maybe because the two channels serve very different physiological function in the nervous(10) and cardiac(52) systems.

VCF data from KCNQ2* channels bearing the S4 charge neutralization mutant R198Q also supports the notion that the fluorescence signal tracks conformational changes of S4 coupled to channel opening. The *de novo* mutation R198Q in KCNQ2 channels has been previously reported to cause infantile spasms with hypsarrhythmia and encephalopathy associated with severe developmental delay(43). Compared with KCNQ2*, KCNQ2*-R198Q channels display left-shifted G(V) and F(V) curves and faster activation and slower deactivation kinetics of current and fluorescence (Figure 5B, C and Figure 5-figure supplement 1A, B), as if the R198Q mutation directly affect S4 movement. Since R198Q seems to ease channel opening by simply altering the voltage sensor, and seemingly not by increasing ionic conductance, this might have potential therapeutic interest as drugs designed to target the S4 in a manner that rightward shifts its voltage dependence could decrease channel opening. Of note, our study did not aim to mechanistically explain how gain-of-function mutations like R198Q cause epilepsy *in vivo*, which would further require the expression of combined wild-type and mutated subunits to mimic the heterozygous state, ideally using inhibitory and excitatory networks of neurons.

Furthermore, fluorescence data from the epilepsy-associated mutation R214W shows a marked separation between the G(V) and F(V) and a seemingly faster fluorescence time course compared to the ionic current time course, suggesting that R214W changes the VSD-PD coupling of KCNQ2*. Previous studies in the related KCNQ1 channel showed that the F351A mutation separated F(V) from G(V)(36), similar to the effect of R214W in KCNQ2 that we show here. Mechanistically, it was postulated that KCNQ1-F351 couples S4 to pore opening possibly through a physical interaction of F351 with residues within the S4-S5 linker such that point mutations like F351A would alter the VSD-PD interactions(36, 37, 45). However, unlike the F351 residue that is localized in the S6 helix (PD) pointing towards the S4-S5 linker of KCNQ1 channels(53), the R214 residue of KCNQ2 lies in the loop region that connects the S4 to the S4-S5 linker(22) (Figure 5-figure supplement 2A). We gained insight into how mutations in this region may alter channel gating from the recent cryo-EM structures of KCNQ1, KCNQ4 and KCNQ2 channels (22, 53, 54). Interestingly, the cryo-EM structure of KCNQ1 and KCNQ4, which captured PIP_2_ bound to these channels, showed that PIP_2_ localizes close to the S4/S4-S5 interface, in a positively charged pocket proposed to be a binding site for the negatively charged PIP_2_. Taking advantage of these findings, we superimposed the existing KCNQ2 structure(22) with homologous structure of KCNQ1 bound to PIP_2_(53) (Figure 5-figure supplement 2B). We noted that residue R214 (together with the adjacent R213 residue) lies very close to PIP_2_, raising the possibility that R214 in the KCNQ2 channel could also be part of the positively charged pocket that coordinates PIP_2_ binding (Figure 5-figure supplement 2B). Previous studies in KCNQ1 and KCNQ3 channels have shown that PIP_2_ directly affects the VSD-PD coupling since PIP_2_ depletion from the membrane impeded pore opening without affecting S4 movement(42, 55). However, the molecular details by which PIP_2_ mediates VSD-PD coupling is still the subject of ongoing debate. Based on our VCF results shown in Figure 5D, E that suggest that R214W mutant would dissociate S4 movement from channel opening, we hypothesize that the positive charge of residue R214 is crucial for PIP_2_ binding. We propose that in KCNQ2, PIP_2_ may act like a molecular “glue” that tightly ties the loop connecting S4 and S4-S5 linker such that during depolarization, the S4 movement effectively pulls S4-S5 away from the pore domain to activate potassium conductance. Therefore, charge-neutralizing mutations at position 214, like the R214W variant studied here, would affect PIP_2_ binding and, thereby, weaken the coupling between the S4 movement and channel opening. This can be interpreted in our allosteric gating model as if the R214W mutation would weaken opening thereby preventing the channel transitioning from closed to open states at negative voltages and only restricting channel opening after multiple voltage sensors are activated at more depolarized voltages (C_4_-to-O_4_ transition, Figure 6F, dashed maroon arrow). Supporting the idea that PIP_2_ may bind to the arginine at position 214, previous studies on KCNQ2 channels bearing the charge neutralizing R214Q or R214W mutations found that the loss of the positive charge, and not changes in residue size, was the main functional effect of these disease-causing mutations as both smaller hydrophilic glutamine and bulkier aromatic tryptophan residues at position 214 favored the resting conformation of S4 segment and, as such, promoted more channel closure(31).

In summary, the results presented in this paper provide a foundation to mechanistically understand the voltage-controlled S4 activation that promotes KCNQ2 channel opening. To date, gating current measurements have not been resolved in KCNQ2 channels, likely due to its slow kinetics. Our allosteric gating scheme, cysteine accessibility, and fluorescence data adds to the existing biophysical and chemical tools to study how KCNQ2 channels open and close the pore in response to changes in the transmembrane voltage. Moreover, the activation scheme proposed here, where a single or multiple voltage sensor movement opens KCNQ2 channel, will lay the groundwork to study molecular mechanisms underlying neurotransmitter-mediated channel inhibition through both PIP_2_ depletion and PKC phosphorylation. This model also provides a mechanistic framework to understand how disease-causing mutations may affect channel gating and how drugs can modulate channel function. Understanding which parameters are affected within our model could provide insight into what region may cause channel dysfunction, as exemplified in the epilepsy-causing uncoupling KCNQ2-R214W mutation.

## Materials and Methods

### Chemicals

[2-(Trimethylammonium)ethyl]methanethiosulfonate Bromide (MTSET) and sodium (2- sulfonatoethyl) methanethiosulfonate (MTSES) were purchased from Toronto Research Chemicals Inc (Downsview, ON, Canada). Alexa Fluor 488 C5-maleimide and Dylight-488-maleimide were purchased from Thermo Fisher Scientific (Waltham, MA, USA). All other chemicals were obtained from Sigma-Aldrich (St. Louis, MO, USA).

### Molecular Biology

The full length human KCNQ2 construct (NCBI Reference Sequence: NP_742105.1; GI: 26051264) was synthesized (GenScript USA, Piscataway, NJ) and ligated between the BamHI and XbaI sites in the multiple cloning site of into the pGEM-HE vector. This vector had been previously modified to contain a T7 promoter and 3’ and 5’ untranslated regions from the Xenopus *β*-globin gene (56). A BglII restriction site (AGATCT) and a Kozak consensus sequence (GCCACC) were added before the start codon (ATG) of the KCNQ2 gene. Point mutations were made in the KCNQ2 gene using the Quikchange XL site-directed Mutagenesis kit (Agilent) according to the manufacturer’s protocol. The correct incorporation of the specific variant was assessed by Sanger sequencing (sequencing by Genewiz LLC, South Plainfield, NJ). The RNA was synthesized in vitro using the mMessage mMachine T7 RNA Transcription Kit (ThermoFisher Scientific) from the linearized cDNA. mRNA (40-50nL) was injected into Xenopus leavis oocytes (purchased from Ecocyte) using a NanojectII nanoinjector (Drummond Scientific) and electrophysiological experiments were performed 2 to 5 days after injection.

### Cysteine accessibility measurements in TEVC recordings

We performed cysteine accessibility to membrane-impermeant thiol methanethiosulfonate (MTS) reagents in two-electrode voltage-clamp (TEVC) recordings as previously described(26, 56). Regular ND96 solution for TEVC contained 96 mM NaCl, 2 mM KCl, 1 mM MgCl_2_, 1.8 mM CaCl_2_ and 5 mM HEPES (pH = 7.5). Stock concentrations of 1-100mM MTS reagents were prepared in distilled water (prechilled to +4 °C) and stored at −20 °C until needed. The MTS-reagents were diluted to the appropriate concentration in ND96 solution just prior to each recording (∼30 s prior to perfusion) and kept on ice for 30 min maximum. We delivered High K+ solution (100mM KCl, 1.8mM CaCl2, 1mM MgCl2, 5mM HEPES, pH 7.5, adjusted with KOH) before each day of experiments (prior to application of MTS-reagents) to check that the rate of wash-in and wash-out of solutions was fast enough to deliver short durations of MTS- reagents to the oocyte (Figure 1-figure supplement 2). A computer-driven valve controlled a home-made perfusion system that allowed for a rapid switching (within 2 s) between ND96 and MTS-reagents during either the open or closed protocol.

We adapted the open and closed state protocols(26) to study the solvent exposure of the substituted cysteines (cys) in S3-S4 and S4 and test whether these cys-residues were exposed in open and/or closed channels using irreversible covalent modification by MTS reagents (Figure 1B). Briefly, cells were held at −80 mV for 1-s before stepping to +20 mV for 12-s, then repolarized for another 12-s to −80 mV (for the open state) or voltages between −80 and −140mV (for the closed state), before stepping to the test potential (+20 mV) to measure the change in currents induced by several 5-s cycles of MTS-reagents (see black rectangles in Figure 1B). We repeat 5-s MTSET application in between 25-s washouts for 8-12 cycles, as shown in the open and closed protocols in Figure 1B. MTS-reagent concentrations were between 1-100 μM. Ionic currents were recorded in TEVC using an OC-725C oocyte clamp (Warner Instruments), low-pass filtered at 1 kHz and sampled at 5 kHz. Microelectrodes were pulled using borosilicate glass to resistances from 0.3 to 0.5 MΩ when filled with 3 M KCl. Voltage clamp data were digitized at 5 kHz (Axon Digidata 1440A; Molecular Devices), collected using pClamp 10 (Axon Instruments). The rate of modification was measured by plotting the change in the current by the MTS reagent (e.g. MTSET) as a function of the exposure to the MTSET (exposure = concentration MTSET [mM] x time [s], measured in (M s)) and fitted with an exponential equation of the form (I(exposure) = I_0_ exp(−exposure/*τ*). We then calculated the second order rate constant from the *τ* values (in M s) as 1/*τ* = *k*_open_ (M^−1^s^−1^) of the MTS reaction. Results are presented as mean ± SEM (n = number of measurements).

### Voltage clamp fluorometry (VCF)

VCF experiments were carried out as previously reported(56). Briefly, aliquots of 50 ng of mRNA coding for KCNQ2 or the KCNQ2 variant RNA were injected into *Xenopus laevis* oocytes. At 2-5 days after injection, oocytes were labeled for 30 min with either 100 μM Alexa-488 maleimide or 100 μM DyLight-488 maleimide (Thermo Fisher Scientific) in high [K^+^] solution (98 mM KCl, 1.8 mM CaCl_2_, 1 mM MgCl_2_, 5 mM HEPES, pH 7.05) at 4 °C, in the dark. The labelled oocytes were then rinsed three-to-five times in dye free ND96 and kept on ice before each recording to prevent internalization of labeled channels. Oocytes were placed into a recording chamber animal pole “up” in ND96 solution (pH 7.5 with NaOH) and electrical measurements were carried out in TEVC using an Axoclamp 900A amplifier (Molecular Devices). Microelectrodes were pulled to resistances from 0.3 to 0.5 MΩ when filled with 3 M KCl. Voltage clamp data were digitized at 5 kHz (Axon Digidata 1550B via a digital Axoclamp 900A commander, Molecular Devices) and collected using pClamp 10 (Axon Instruments). Fluorescence recordings were performed using an Olympus BX51WI upright microscope. Light was focused on the top of the oocyte through a 20× water immersion objective after being passed through an Oregon green filter cube (41026; Chroma).

Fluorescence signals were focused on a photodiode and amplified with an Axopatch 200B patch clamp amplifier (Axon Instruments). Fluorescence signals were low-pass Bessel-filtered (Frequency Devices) at 100–200 Hz, digitized at 1 kHz, and recorded using pClamp 10. When needed, we added 100 μM LaCl_3_ to the batch solution to block endogenous hyperpolarization-activated currents. At this concentration, La^3+^ did not affect G(V) or F(V) curves from KCNQ2 channels.

### Rb^+^/K^+^ permeability

Experiments were recorded under TEVC using 100 mM Na^+^ solution (96 mM NaCl, 4 mM KCl, 1.8 mM CaCl2, 1 mM MgCl2, 5 mM HEPES, 100 mM K^+^ solution (100 mM KCl, 1.8 mM CaCl2, 1 mM MgCl2, 5 mM HEPES or 100 mM Rb^+^ solution (96 mM RbCl, 4 mM KCl, 1.8 mM CaCl2, 1 mM MgCl2, 5 mM HEPES); pH = 7.5. Rb^+^/K^+^ permeability ratios were determined using inward tail current measured at −60 mV following the 2-s activation pulse at +60 mV. Inward tail currents were determined following normalization of outward current to calculate Rb^+^/K^+^ ratio.

### Modelling

Fluorescence and current from wt-KCNQ2 and mutated KCNQ2 channel models were simulated using Berkeley Madonna software (Berkeley, CA, USA), similar to the allosteric model initially proposed for KCNQ1 channels(36). In our KCNQ2 model, rate constants for each transition were of the form k_i_ = k_o_ x exp(-z_i_(V)/*k*T), where k_i_ and z_i_ were determined from data in Figure 3 and Figure 6 (see all parameters in supplement Table 2). R, T and F have their usual thermodynamic meaning.

The homology model of KCNQ2 channels with S4 in the resting (down) state was created using the Swiss-model program (https://swissmodel.expasy.org/) with the model of KCNQ1 in the resting state(57), as template. All images were created in UCSF ChimeraX, version 1.1 (2020-10-07).

### Electrophysiology data Analysis

Data were analyzed with Clampfit 10 (Axon Instruments, Inc., Sunnyvale, CA), OriginPro 2021b (OriginLabs Northampton, MA), and Corel-DRAW Graphics Suite 2021 software. To determine the ionic conductance established by a given test voltage, a test voltage pulse was followed by a step to the fixed voltage of –40 mV (tail), and current was recorded following the step. To estimate the conductance g(V) activated at the end of the test pulse to voltage V, the current flowing after the hook was exponentially extrapolated to the time of the step and divided by the offset between –40 mV and the reversal potential. The conductance g(V) associated with different test voltages V in a given experiment was fitted by the relation:

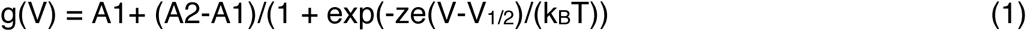

where A1 and A2 are conductances that would be approached at extreme negative or positive voltages, respectively, V_1/2_ the voltage that activates the conductance (A1+A2)/2, and z is an apparent valency describing the voltage sensitivity of activation (e is the electron charge, k_B_ the Boltzmann constant, and T the absolute temperature). Due to the generally different numbers of expressed channels in different oocytes, we compare normalized conductance, G(V):

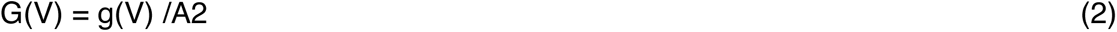

Fluorescence signals were corrected for bleaching and time-averaged over 10-40 ms intervals for analysis. The voltage dependence of fluorescence f(V) was analyzed and normalized (F(V)) using relations analogous to those for conductance (equations. 1 and 2).

### Statistics

All experiments were repeated 4 or more times from at least three batches of oocytes. Pairwise comparisons were achieved using Student’s t test or ANOVA with a Tukey′s test. Data are represented as mean ± SEM (standard error of mean) and “n” represents the number of experiments.

## Acknowledgements

We thank Drs. Derek Dykxhoorn and Hans Peter Larsson for helpful comments on the manuscript. This work was supported by the National Institutes of Health (1R01NS110847) to Rene Barro-Soria.

## Supplementary Information

**Figure 1-figure supplement 1.**
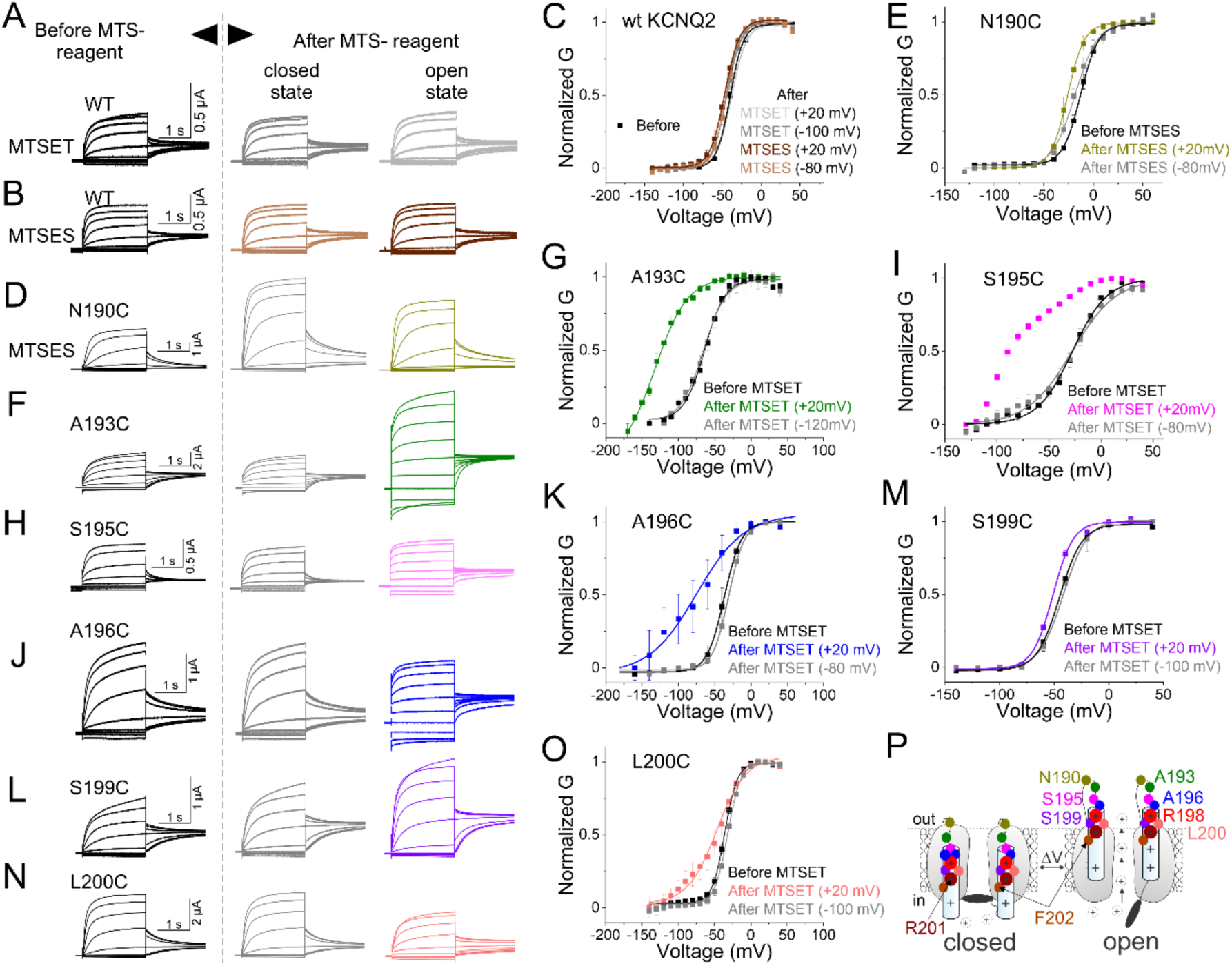
State-dependent modification of S4 residues by external MTS-reagents consistent with outward S4 motion. (A-H) Currents from oocytes expressing (A and B) wt, (D) N190C, (F) A193C, (H) S195C, (J) A196C, (L) S199C, and (N) L200C channels in response to 20-mV voltage steps from –140 mV to +40 mV (left) before MTSET, (middle) after several 5-s applications of MTSET at –80 mV, and (right) after several 5-s applications of MTSET at +20 mV. We repeat 5-s MTSET (or MTSES in B and D) application in between 25-s washouts for 8-19 cycles, as shown in the open and closed protocols in Figure 1B. We used MTSET concentrations ranging from 1 to 100mM, respectively. (C, E, G, I, K, M and O) Normalized G(V) relations of recordings from panels (A and B), (D), (F), (H), (J), (L), and (N), respectively, before (black) and after MTSET application in the closed (–80 mV, gray) and open (+ 20mV, color coded) states. mean ± SEM, n=3-11. Lines represent the fitted theoretical voltage dependencies (equations1 and 2). (P) Cartoon representing the voltage-dependent cysteine accessibility data from all residues assayed. Unlike residue N190 (dark yellow) that is modified by MTSET in both closed and open states (always exposed), residue F202 (brown) remains unmodified in both closed and open states (buried in the membrane). A stretch of 8-9 amino acids (193 to 200-201) moves from a membrane-buried position in the closed state to the extracellular solution during channel opening. Note that because R201C produces voltage-independent channels, we cannot test the state-dependent modification of MTS-reagents. The dashed line indicates the proposed outer lipid bilayer boundary. Only two subunits of the tetrameric channel are shown.

**Figure1-figure supplement 2.**
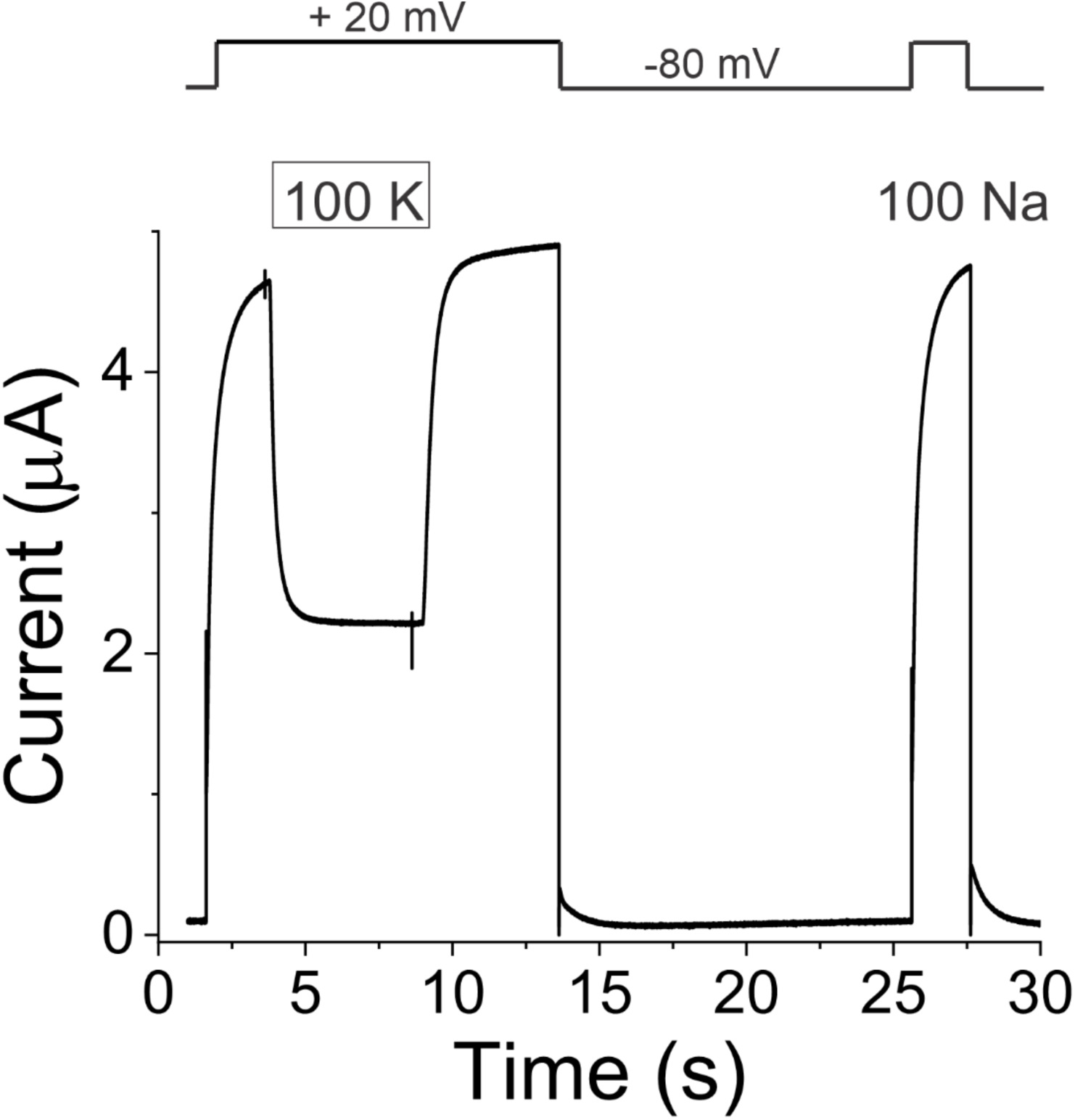
Fast perfusion system delivers 5-s applications of external solution exchange to whole oocytes. Representative time course of solution exchange from 100 mM NaCl (Na) to 100 mM KCl (K). Currents from KCNQ2 channels in response to a +20-mV pulse from a holding potential of –80 mV followed by a tail potential of –80 mV. Extracellular solution was ND96 (100 mM NaCl) except for the 5-s application for which the 100 mM NaCl was exchanged for 100 mM KCl. Shown are 3 cycles of solution exchanging as shown in the protocol (top). The application of 100 mM KCl quickly reduces (*τ* = 0.21 ± 3.6 s) the outward currents and the reintroduction of the 100 mM NaCl quickly (*τ* = 0.32 ± 1.5 s) restores the currents.

**Figure 1-figure supplement 3.**
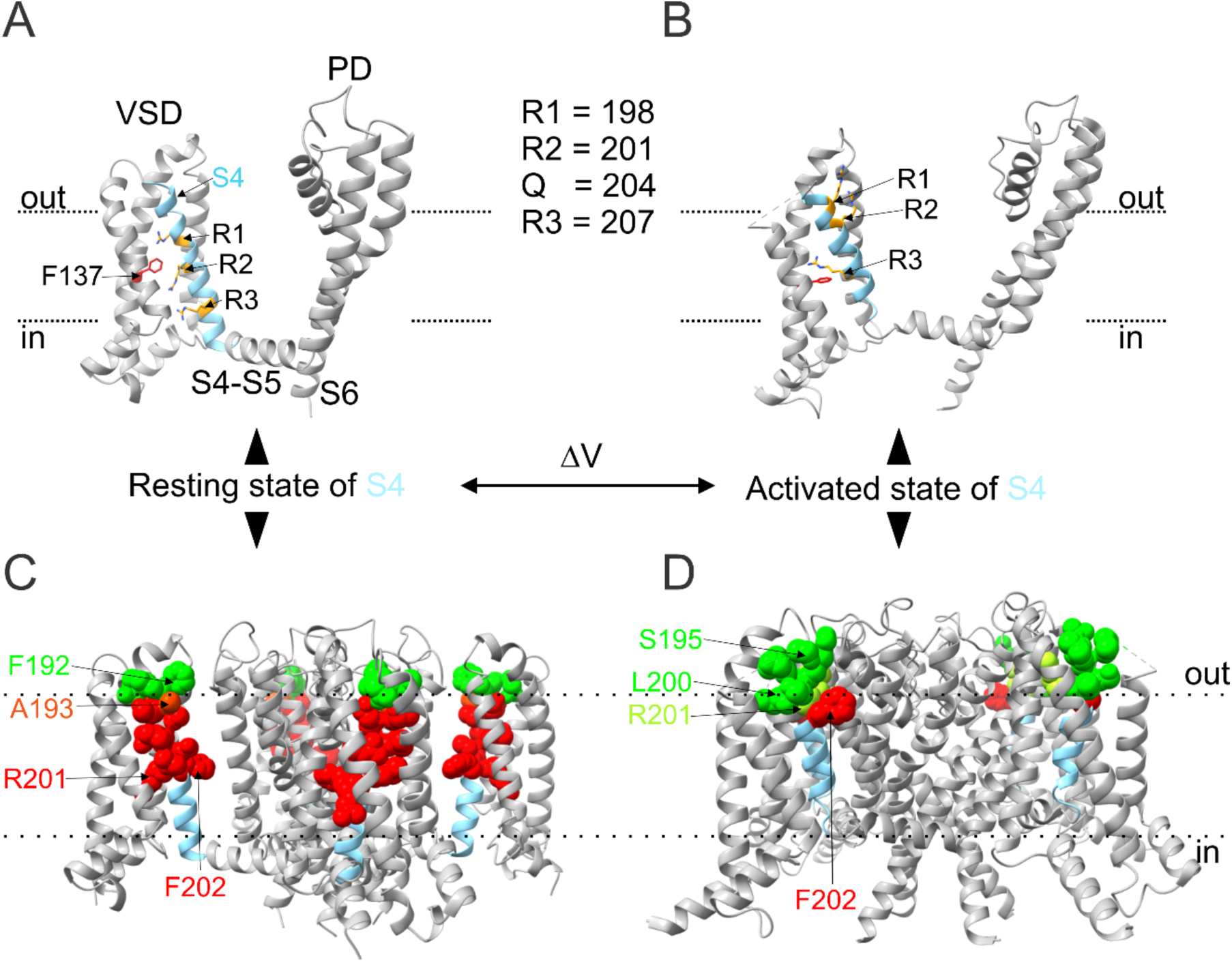
Proposed molecular motions of S4 residues in KCNQ2 channels. (A and C) KCNQ2 homology model in the closed resting state (S4 down) and (B and D) Cryo-EM structure of KCNQ2 channel in the activated state of S4 (up) and closed pore. (A) In the resting state, R1 and R2 in S4 (cyan) localize above and below the gating charge transfer center F137 (red stick), respectively. (B) Upon S4 activation, R1 and R2 move about three helical turns outward from F137 into a position close to or within the extracellular space. One subunit is shown as ribbons and key amino acid residues as sticks. VSD: voltage sensing domain; PD: pore domain (PD). (C-D) Proposed molecular motions of S4 residues from (C) resting to (D) activated states from cysteine accessibility data. A buried (red spheres) stretch of 8-9 amino acids (193 to 200-201) in the resting state (C) becomes exposed to the extracellular space (green spheres) in the activated state (D). The four subunits are shown as ribbons and buried and extracellularly exposed residues in the S4 are shown as red and green spheres, respectively. Dotted lines indicate the proposed inner and outer lipid bilayer boundary. PDB code for KCNQ2: 7CR0. All images were generated using UCSF ChimeraX, version 1.1 (2020-10-07).

**Figure 2-figure supplement 1.**
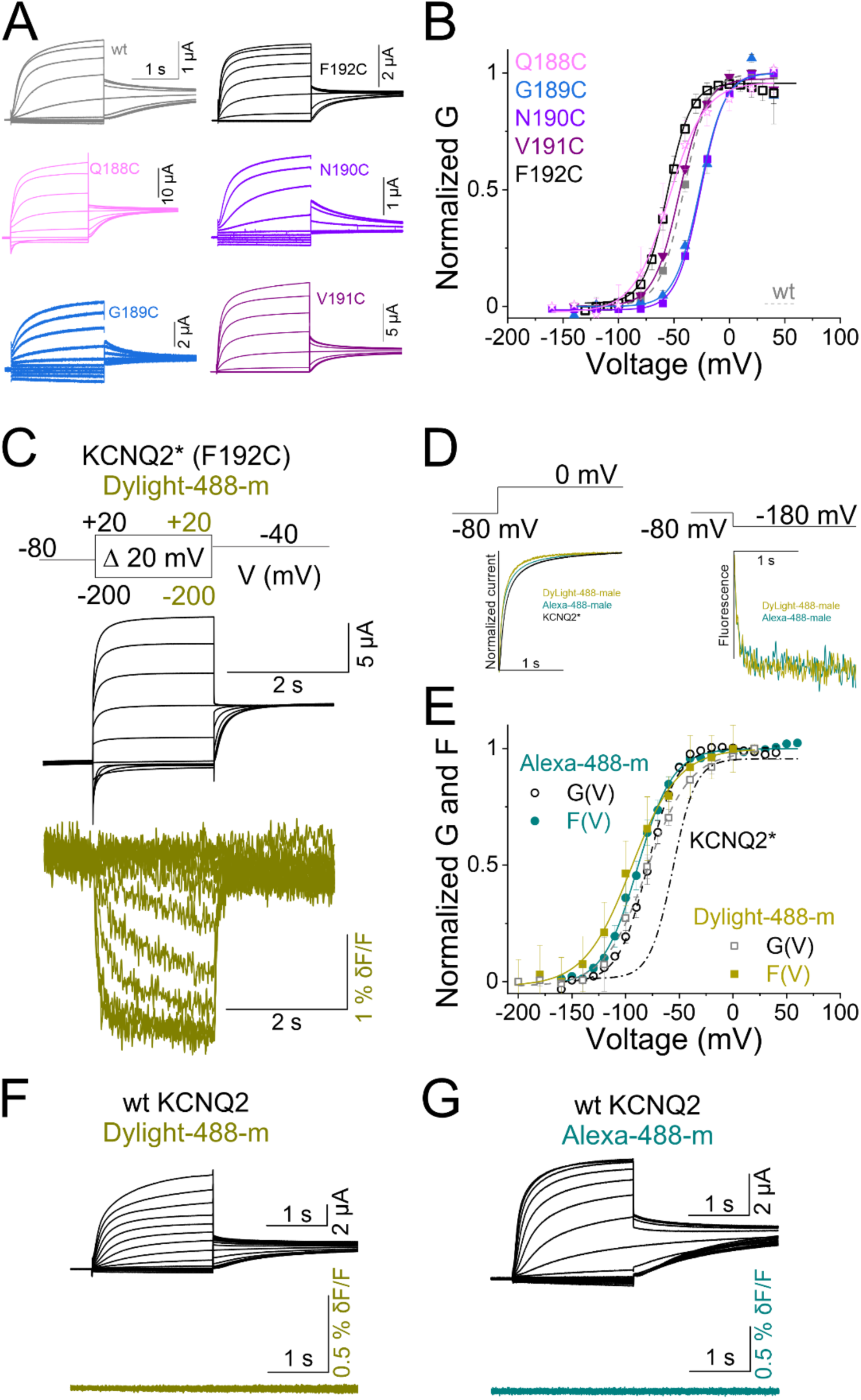
Cysteine-scan mutagenesis of S3-S4 linker identifies F192C as the ideal position for fluorophore labeling. (A) Currents from oocytes expressing a series of cysteine mutants in the S3-S4 linker of KCNQ2 channel. Cells are held at −80 mV and stepped to potentials between −140 mV and +40 mV in 20-mV steps for 2 s followed by a tail to −40 mV. (B) Normalized G(V) curves from cysteine mutations from (A). Dashed line represents wt KCNQ2. Data are mean ± SEM, (*n= 3-8*). (C) Representative current (black) and fluorescence (dark yellow) traces from Dylight-488-labeled KCNQ2-F192C channels (KCNQ2*) for the indicated voltage protocol (top). (D) Comparison of activation kinetics of (left) current and (right) fluorescence signals from (dark yellow) Dylight-488- and (cyan) Alexa-488- labeled KCNQ2* channels in response to the voltage protocol shown. For comparison, the activation kinetics of current (black solid line) curve of KCNQ2* is shown. (E) Normalized G(V) (open symbols and dashed lines from a Boltzmann fit) and F(V) (closed symbols and solid lines from a Boltzmann fit) curves from (dark yellow) Dylight-488- and (cyan) Alexa-488- labeled KCNQ2* channels. For comparison, the G(V) (dashed line) curve of KCNQ2* is shown. Data are mean ± SEM, (*n= 8-11*). The midpoints of activation for the fits are shown in supplement Table 1. (F-G) Representative (top panels) current and (bottom panels) fluorescence traces from (F) Dylight-488- and (G) Alexa-488-incubated wt KCNQ2 channels.

**Figure 5-figure supplement 1.**
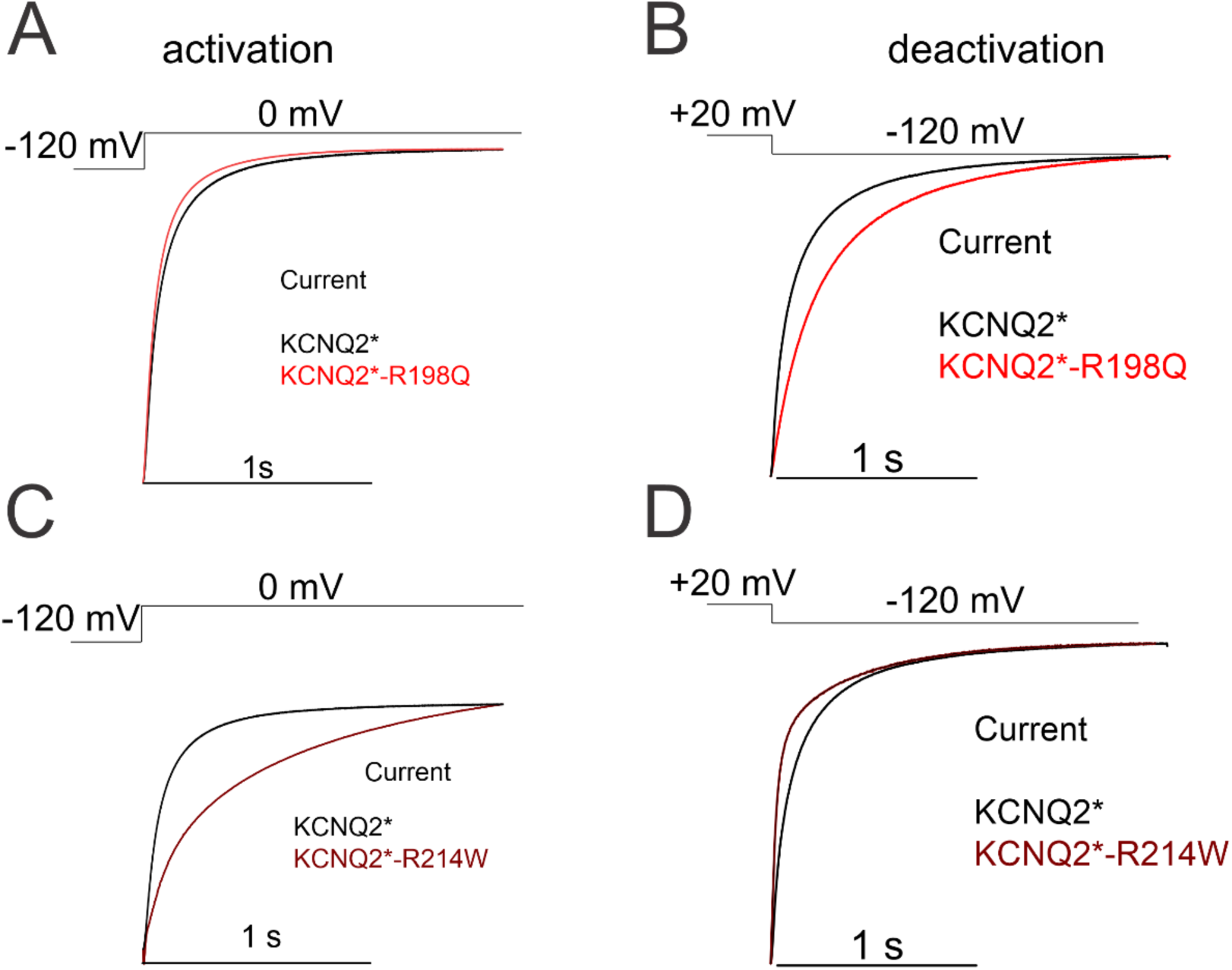
The kinetics of epilepsy-causing mutations- R198Q and R214W differ from wt channel. Normalized time courses of current (A and C) activation and (B and D) deactivation from (red) R198Q and (maroon) R214W mutated channels in response to the indicated voltage step for 2 s from the showed holding potentials.

**Figure 5-figure supplement 2.**
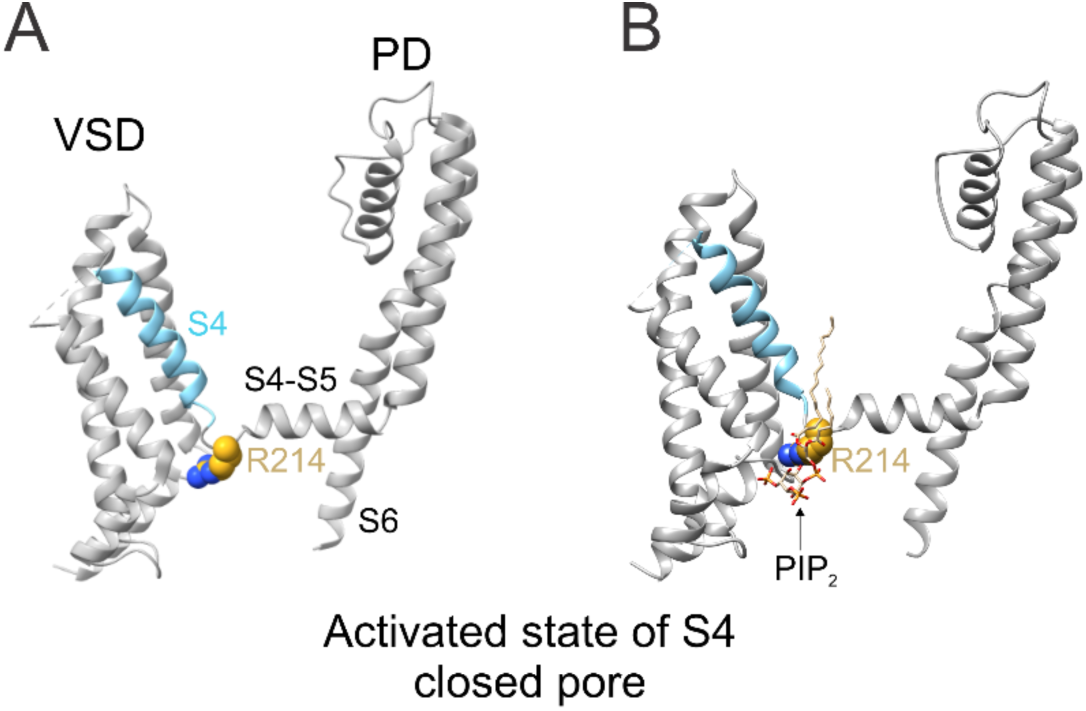
PIP2 tightly joints the loop connecting S4 and S4-S5 linker to facilitate channel opening. Cryo-EM structure of KCNQ2 channel in the activated state (S4 (up) and closed pore showing (A) the localization of residue R214 (sphere) in the loop connecting S4 (cyan) and S4-S5 linker, and (B) the predicted position of PIP2 relative to R214 within the S4/S4-S5 interface of KCNQ2 channels. PDB code for KCNQ2: 7CR0. All images were generated using UCSF ChimeraX, version 1.1 (2020-10-07).

**Figure 6-figure supplement 1.**
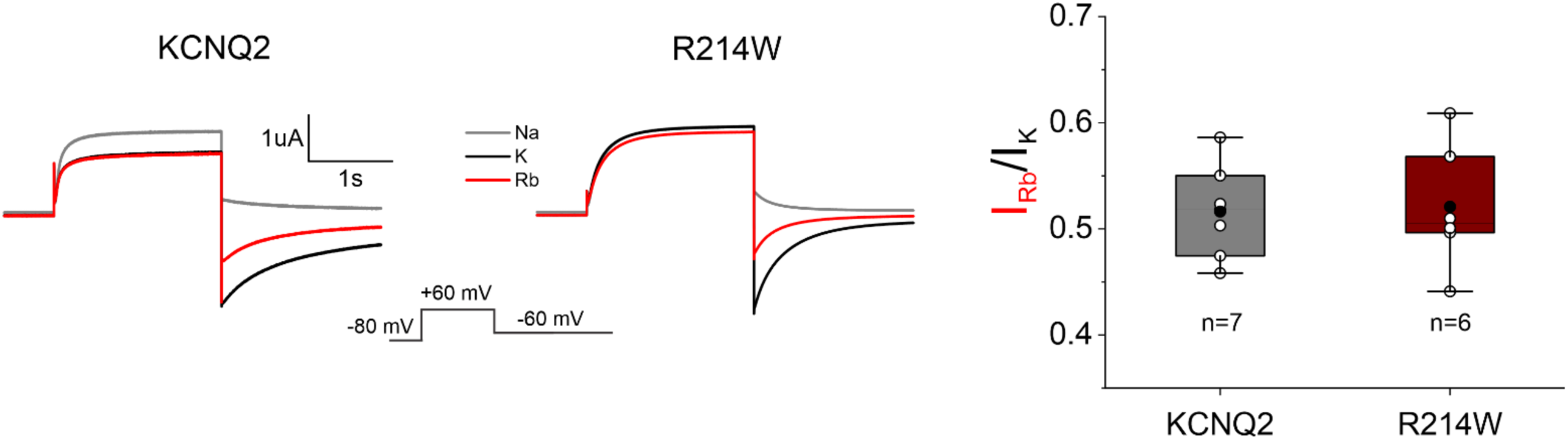
The permeation properties of KCNQ2 channels. (A) Current traces from oocytes expressing (left) wt-KCNQ2 and (right) KCNQ2-R214W channels under high external K^+^ (black), Rb^+^ (red), and Na^+^ (gray) concentration. Cells are held at −80 mV and stepped to +60 mV followed by a step to −60 mV. (B) Comparison of measured tail Rb^+^/K^+^ ratio from the KCNQ2 and R214W channels. Data are shown as mean ± SEM, p=0.87

**Supplement Table 1.** Biophysical properties of wild type and mutant KCNQ2 channels. V_1/2_ and F_1/2_ of activation; V_1/2_ of state dependent MTS modification, and the second order rate constant of KCNQ2 channels. Data are mean ± SEM, n = number of cells.

| Constructs | n | $V_{1/2}$ (mV) | n | $F_{1/2}$ (mV) | n | $V_{1/2}$ (mV)<br>open state<br>MTSET | n | $V_{1/2}$ (mV)<br>closed<br>state<br>MTSET | Second<br>order<br>rate<br>constant<br>($M^{-1}s^{-1}$)<br>(open<br>state) | Second<br>order<br>rate<br>constant<br>($M^{-1}s^{-1}$)<br>(closed<br>state) |
| --- | --- | --- | --- | --- | --- | --- | --- | --- | --- | --- |
| wt-KCNQ2<br>(+MTSET) | 8 | $-39.8 \pm 0.5$ | - | - | 5 | $-44.2 \pm 0.5$ | 3 | $-42.1 \pm 0.4$ | - | - |
| wt-KCNQ2<br>(+MTSES) | 3 | $-45.4 \pm 1$ | - | - | 3 | $-47.3 \pm 0.7$ | 3 | $-44.9 \pm 0.6$ | - | - |
| Q188C | 5 | $-54.3 \pm 2.1$ | - | - | - | - | - | - | - | - |
| G189C | 3 | $-44.47 \pm 7.5$ | - | - | - | - | - | - | - | - |
| N190C<br>(+MTSES) | 5 | $-12.9 \pm 0.5$ | - | - | 3 | $-26 \pm 0.3$ | 3 | $-18.5 \pm 0.8$ | - | - |
| N190C | 6 | $-26.1 \pm 0.1$ | - | - | 4 | $-45 \pm 1.3$ | 3 | $-29.77 \pm 0.1$ | - | - |
| V191C | 4 | $-45.0 \pm 1.2$ | - | - | - | - | - | - | - | - |
| F192C | 8 | $-50.1 \pm 2.2$ | - | - | - | - | - | - | - | - |
| A193C | 11 | $-63.9 \pm 1.4$ | - | - | 7 | $-112.2 \pm 2.4$ | 3 | $-65.7 \pm 1.7$ | $11800 \pm 2.6$<br>at +20-<br>mV | $2080.3 \pm 1.9$<br>at -100-mV |
| S195C | 3 | $-24.1 \pm 4.7$ | - | - | 3 | $-142.6 \pm 19.7$ | 3 | $-27.8 \pm 2.4$ | $11350 \pm 0.01$ | $38.6 \pm 1.1$ |
| A196C | 6 | $-36.8 \pm 0.5$ | - | - | 3 | $-75.1 \pm 7$ | 2 | $-30.8 \pm 0.7$ | $1400 \pm 1.1$ | $94.3 \pm 9.6$ |
| R198C | 8 | $-32.01 \pm 0.3$ | - | - | 3 | $-87.5 \pm 1.5$ | 3 | $-29.9 \pm 0.6$ | $3230 \pm 2.05$ | $7.03 \pm 3.2$ |
| S199C | 10 | $-44.3 \pm 0.6$ | - | - | 3 | $-51.3 \pm 0.8$ | 3 | $-42.3 \pm 1$ | $334.6 \pm 3.2$ | $11.9 \pm 1.7$ |
| L200C | 3 | $-31.9 \pm 0.5$ | - | - | 3 | $-41 \pm 2.6$ | 3 | $-31.7 \pm 0.7$ | $28.2 \pm 5.8$ | $4.4 \pm 7.9$ |
| R201C | 4 | Voltage<br>independent | - | - | - | - | - | - | - | - |
| F202C | 5 | $-33.5 \pm 0.4$ | - | - | 4 | $-31 \pm 2.6$ | 4 | $-31.3 \pm 2.3$ | - | - |
| F192C/Alexa4<br>88 maleimide<br>(KCNQ2*) | 8 | $-77.1 \pm 2.7$ | 7 | $-87.1 \pm 3.9$ | - | - | - | - | - | - |
| F192C/<br>Dylight-488<br>maleimide | 11 | $-79.9 \pm 1.4$ | 8 | $-94.7 \pm 1.8$ | - | - | - | - | - | - |
| KCNQ2*-<br>R198Q | 10 | $-110.3 \pm 3.5$ | 4 | $-119.8 \pm 4.2$ | - | - | - | - | - | - |
| KCNQ2*-<br>R214W | 8 | $-17.1 \pm 0.9$ | 7 | $-77 \pm 0.6$ | - | - | - | - | - | - |

**Supplement Table 2.** Parameters for model in Figure 6. L represents an allosteric coupling factor associated with the backward rates in the model of Figure 6F. The gating charges z associated with each transition were determined from fits of the data in Figure 3, where ki = ko x exp(-zi(V)/kT).

| Parameters |  |  |
| --- | --- | --- |
| Rate | *KCNQ2* | *R214W* |
| $z\alpha$ | 0.43 ($e0$) | 0.43 ($e0$) |
| $z\beta$ | 1.18 ($e0$) | 1.18 ($e0$) |
| $\gamma$ | 0.17 s<sup>-1</sup> | 0.003 s<sup>-1</sup> |
| $\delta$ | 9.97 s<sup>-1</sup> | 9.97 s<sup>-1</sup> |
| L | 1.5 | 7.4 |
| f | 3.11 | 2500 |
| $V_{1/2}$ | -72 mV | -37 mV |

## References

1. Jespersen T, Grunnet M, Olesen SP. The KCNQ1 potassium channel: from gene to physiological function. Physiology (Bethesda). 2005;20:408–16.

2. Jentsch TJ. Neuronal KCNQ potassium channels: physiology and role in disease. Nature Reviews Neuroscience. 2000;1(1):21–30.

3. Abbott GW, Pitt GS. Ion channels under the sun. FASEB J. 2014;28(5):1957–62.

4. Hille B. Ion Channels of Excitable Membranes. 3rd ed. Sunderland, MA: Sinauer Associates, INC; 2001.

5. Brown DA, Adams PR. Muscarinic suppression of a novel voltage-sensitive K+ current in a vertebrate neurone. Nature. 1980;283(5748):673–6.

6. Halliwell JV, Adams PR. Voltage-clamp analysis of muscarinic excitation in hippocampal neurons. Brain Res. 1982;250(1):71–92.

7. Wang HS, Pan Z, Shi W, Brown BS, Wymore RS, Cohen IS, et al. KCNQ2 and KCNQ3 potassium channel subunits: molecular correlates of the M-channel. 1998;282(5395):1890–3.

8. Schroeder BC, Hechenberger M, Weinreich F, Kubisch C, Jentsch TJ. KCNQ5, a novel potassium channel broadly expressed in brain, mediates M-type currents. Journal of Biological Chemistry. 2000.

9. Soh H, Pant R, LoTurco JJ, Tzingounis AV. Conditional deletions of epilepsy-associated KCNQ2 and KCNQ3 channels from cerebral cortex cause differential effects on neuronal excitability. J Neurosci. 2014;34(15):5311–21.

10. Delmas P, Brown DA. Pathways modulating neural KCNQ/M (Kv7) potassium channels. Nat Rev Neurosci. 2005;6(11):850–62.

11. Adams PR, Brown DA, Constanti A. Pharmacological inhibition of the M-current. J Physiol. 1982;332:223–62.

12. Adams PR, Jones SW, Pennefather P, Brown DA, Koch C, Lancaster B. Slow synaptic transmission in frog sympathetic ganglia. J Exp Biol. 1986;124:259–85.

13. Greene DL, Hoshi N. Modulation of Kv7 channels and excitability in the brain. Cell Mol Life Sci. 2017;74(3):495–508.

14. Geisheker MR, Heymann G, Wang T, Coe BP, Turner TN, Stessman HAF, et al. Hotspots of missense mutation identify neurodevelopmental disorder genes and functional domains. Nat Neurosci. 2017;20(8):1043–51.

15. Weckhuysen S, Mandelstam S, Suls A, Audenaert D, Deconinck T, Claes LR, et al. KCNQ2 encephalopathy: emerging phenotype of a neonatal epileptic encephalopathy. Ann Neurol. 2012;71(1):15–25.

16. Weckhuysen S, Ivanovic V, Hendrickx R, Van Coster R, Hjalgrim H, Moller RS, et al. Extending the KCNQ2 encephalopathy spectrum: clinical and neuroimaging findings in 17 patients. Neurology. 2013;81(19):1697–703.

17. Orhan G, Bock M, Schepers D, Ilina EI, Reichel SN, Loffler H, et al. Dominant-negative effects of KCNQ2 mutations are associated with epileptic encephalopathy. Ann Neurol. 2014;75(3):382–94.

18. Saitsu H, Kato M, Koide A, Goto T, Fujita T, Nishiyama K, et al. Whole exome sequencing identifies KCNQ2 mutations in Ohtahara syndrome. Ann Neurol. 2012;72(2):298–300.

19. Kato M, Yamagata T, Kubota M, Arai H, Yamashita S, Nakagawa T, et al. Clinical spectrum of early onset epileptic encephalopathies caused by KCNQ2 mutation. Epilepsia. 2013;54(7):1282–7.

20. Rauch A, Wieczorek D, Graf E, Wieland T, Endele S, Schwarzmayr T, et al. Range of genetic mutations associated with severe non-syndromic sporadic intellectual disability: an exome sequencing study. Lancet. 2012;380(9854):1674–82.

21. Cornet MC, Sands TT, Cilio MR. Neonatal epilepsies: Clinical management. Semin Fetal Neonatal Med. 2018;23(3):204–12.

22. Li X, Zhang Q, Guo P, Fu J, Mei L, Lv D, et al. Molecular basis for ligand activation of the human KCNQ2 channel. Cell Res. 2021;31(1):52–61.

23. Long SB, Campbell EB, Mackinnon R. Crystal structure of a mammalian voltage-dependent Shaker family K+ channel. Science. 2005;309(5736):897–903.

24. del Camino D, Yellen G. Tight steric closure at the intracellular activation gate of a voltage-gated K(+) channel. Neuron. 2001;32(4):649–56.

25. Mannuzzu LM, Moronne MM, Isacoff EY. Direct physical measure of conformational rearrangement underlying potassium channel gating. Science. 1996;271(5246):213–6.

26. Larsson HP, Baker OS, Dhillon DS, Isacoff EY. Transmembrane movement of the shaker K+ channel S4. Neuron. 1996;16(2):387–97.

27. Seoh SA, Sigg D, Papazian DM, Bezanilla F. Voltage-sensing residues in the S2 and S4 segments of the Shaker K+ channel. Neuron. 1996;16(6):1159–67.

28. Aggarwal SK, MacKinnon R. Contribution of the S4 segment to gating charge in the Shaker K+ channel. Neuron. 1996;16(6):1169–77.

29. Miceli F, Vargas E, Bezanilla F, Taglialatela M. Gating currents from Kv7 channels carrying neuronal hyperexcitability mutations in the voltage-sensing domain. Biophys J. 2012;102(6):1372–82.

30. Soldovieri MV, Ambrosino P, Mosca I, Miceli F, Franco C, Canzoniero LMT, et al. Epileptic Encephalopathy In A Patient With A Novel Variant In The Kv7.2 S2 Transmembrane Segment: Clinical, Genetic, and Functional Features. Int J Mol Sci. 2019;20(14).

31. Miceli F, Soldovieri MV, Hernandez CC, Shapiro MS, Annunziato L, Taglialatela M. Gating consequences of charge neutralization of arginine residues in the S4 segment of K(v)7.2, an epilepsy-linked K+ channel subunit. Biophys J. 2008;95(5):2254–64.

32. Gourgy-Hacohen O, Kornilov P, Pittel I, Peretz A, Attali B, Paas Y. Capturing distinct KCNQ2 channel resting states by metal ion bridges in the voltage-sensor domain. J Gen Physiol. 2014;144(6):513–27.

33. Nerbonne JM, Kass RS. Molecular physiology of cardiac repolarization. Physiol Rev. 2005;85(4):1205–53.

34. Zagotta WN, Hoshi T, Aldrich RW. Shaker potassium channel gating. III: Evaluation of kinetic models for activation. Journal of General Physiology. 1994;103(2):321–62.

35. Horrigan FT, Cui J, Aldrich RW. Allosteric voltage gating of potassium channels I. Mslo ionic currents in the absence of Ca(2+). J Gen Physiol. 1999;114(2):277–304.

36. Osteen JD, Gonzalez C, Sampson KJ, Iyer V, Rebolledo S, Larsson HP, et al. KCNE1 alters the voltage sensor movements necessary to open the KCNQ1 channel gate. Proceedings of the National Academy of Sciences of the United States of America. 2010;107(52):22710–5.

37. Zaydman MA, Kasimova MA, McFarland K, Beller Z, Hou P, Kinser HE, et al. Domain-domain interactions determine the gating, permeation, pharmacology, and subunit modulation of the IKs ion channel. Elife. 2014;3:e03606.

38. Hou P, Eldstrom J, Shi J, Zhong L, McFarland K, Gao Y, et al. Inactivation of KCNQ1 potassium channels reveals dynamic coupling between voltage sensing and pore opening. Nat Commun. 2017;8(1):1730.

39. Hou P, Kang PW, Kongmeneck AD, Yang ND, Liu Y, Shi J, et al. Two-stage electro-mechanical coupling of a KV channel in voltage-dependent activation. Nat Commun. 2020;11(1):676.

40. Bell DC, Yao H, Saenger RC, Riley JH, Siegelbaum SA. Changes in local S4 environment provide a voltage-sensing mechanism for mammalian hyperpolarization-activated HCN channels. The Journal of general physiology. 2004;123(1):5–19.

41. Lakowicz JR. Principles of Fluorescence Spectroscopy. New York: Springer; 2006.

42. Kim RY, Pless SA, Kurata HT. PIP2 mediates functional coupling and pharmacology of neuronal KCNQ channels. Proc Natl Acad Sci U S A. 2017;114(45):E9702–e11.

43. Millichap JJ, Miceli F, De Maria M, Keator C, Joshi N, Tran B, et al. Infantile spasms and encephalopathy without preceding neonatal seizures caused by KCNQ2 R198Q, a gain-of-function variant. Epilepsia. 2017;58(1):e10–e5.

44. Castaldo P, del Giudice EM, Coppola G, Pascotto A, Annunziato L, Taglialatela M. Benign familial neonatal convulsions caused by altered gating of KCNQ2/KCNQ3 potassium channels. J Neurosci. 2002;22(2):RC199.

45. Taylor KC, Kang PW, Hou P, Yang ND, Kuenze G, Smith JA, et al. Structure and physiological function of the human KCNQ1 channel voltage sensor intermediate state. Elife. 2020;9.

46. Pathak M, Kurtz L, Tombola F, Isacoff E. The cooperative voltage sensor motion that gates a potassium channel. J Gen Physiol. 2005;125(1):57–69.

47. Schoppa NE, McCormack K, Tanouye MA, Sigworth FJ. The size of gating charge in wild-type and mutant Shaker potassium channels. Science. 1992;255(5052):1712–5.

48. Zagotta WN, Hoshi T, Dittman J, Aldrich RW. Shaker potassium channel gating. II: Transitions in the activation pathway. J Gen Physiol. 1994;103(2):279–319.

49. Barro-Soria R, Rebolledo S, Liin SI, Perez ME, Sampson KJ, Kass RS, et al. KCNE1 divides the voltage sensor movement in KCNQ1/KCNE1 channels into two steps. Nat Commun. 2014;5:3750.

50. Hodgkin AL, Huxley AF. A quantitative description of membrane current and its application to conduction and excitation in nerve. J Physiol. 1952;117(4):500–44.

51. Osteen JD, Barro-Soria R, Robey S, Sampson KJ, Kass RS, Larsson HP. Allosteric gating mechanism underlies the flexible gating of KCNQ1 potassium channels. Proceedings of the National Academy of Sciences of the United States of America. 2012;109(18):7103–8.

52. Wang Q, Curran ME, Splawski I, Burn TC, Millholland JM, VanRaay TJ, et al. Positional cloning of a novel potassium channel gene: KVLQT1 mutations cause cardiac arrhythmias. Nat Genet. 1996;12(1):17–23.

53. Sun J, MacKinnon R. Structural Basis of Human KCNQ1 Modulation and Gating. Cell. 2020;180(2):340–7 e9.

54. Li T, Wu K, Yue Z, Wang Y, Zhang F, Shen H. Structural Basis for the Modulation of Human KCNQ4 by Small-Molecule Drugs. Mol Cell. 2021;81(1):25–37 e4.

55. Zaydman MA, Silva JR, Delaloye K, Li Y, Liang H, Larsson HP, et al. Kv7.1 ion channels require a lipid to couple voltage sensing to pore opening. Proc Natl Acad Sci U S A. 2013;110(32):13180–5.

56. Barro-Soria R. Epilepsy-associated mutations in the voltage sensor of KCNQ3 affect voltage dependence of channel opening. J Gen Physiol. 2018.

57. Kuenze G, Duran AM, Woods H, Brewer KR, McDonald EF, Vanoye CG, et al. Upgraded molecular models of the human KCNQ1 potassium channel. PLoS One. 2019;14(9):e0220415.

